# Diversity, functional classification and genotyping of SHV β-lactamases in *Klebsiella pneumoniae*

**DOI:** 10.1101/2024.04.05.587953

**Authors:** Kara K. Tsang, Margaret M.C. Lam, Ryan R. Wick, Kelly L. Wyres, Michael Bachman, Stephen Baker, Katherine Barry, Sylvain Brisse, Susana Campino, Alexandra Chiaverini, Daniela Maria Cirillo, Taane Clark, Jukka Corander, Marta Corbella, Alessandra Cornacchia, Aline Cuénod, Nicola D’Alterio, Federico Di Marco, Pilar Donado-Godoy, Adrian Egli, Refath Farzana, Edward J. Feil, Aasmund Fostervold, Claire L. Gorrie, Yukino Gütlin, Brekhna Hassan, Marit Andrea Klokkhammer Hetland, Le Nguyen Minh Hoa, Le Thi Hoi, Benjamin Howden, Odion O. Ikhimiukor, Adam W. J. Jenney, Håkon Kaspersen, Fahad Khokhar, Thongpan Leangapichart, Małgorzata Ligowska-Marzęta, Iren Høyland Löhr, Scott W. Long, Amy J. Mathers, Andrew G. McArthur, Geetha Nagaraj, Anderson O. Oaikhena, Iruka N. Okeke, João Perdigão, Hardik Parikh, My H. Pham, Francesco Pomilio, Niclas Raffelsberger, Andriniaina Rakotondrasoa, K L Ravi Kumar, Leah W. Roberts, Carla Rodrigues, Ørjan Samuelsen, Kirsty Sands, Davide Sassera, Helena Seth-Smith, Varun Shamanna, Norelle L. Sherry, Sonia Sia, Anton Spadar, Nicole Stoesser, Marianne Sunde, Arnfinn Sundsfjord, Pham Ngoc Thach, Nick Thomson, Harry A. Thorpe, Estée Torok, Van Dinh Trang, Nguyen Vu Trung, Jay Vornhagen, Timothy Walsh, Ben Warne, Hayley Wilson, Gerard D. Wright, Kathryn E. Holt, KlebNET AMR Genotype-Phenotype Group

**Affiliations:** Department of Infection Biology, Faculty of Infectious and Tropical Diseases, London School of Hygiene & Tropical Medicine, London WC1E 7HT, UK; Department of Infectious Diseases, Central Clinical School, Monash University, Melbourne, Victoria 3004, Australia; Department of Microbiology and Immunology, The Peter Doherty Institute for Infection and Immunity, University of Melbourne, Melbourne, Victoria, Australia; University of Michigan; University of Cambridge; University of Virginia; Institut Pasteur, Université Paris Cité, Biodiversity and Epidemiology of Bacterial Pathogens, Paris, France; Istituto Zooprofilattico Sperimentale, dell’Abruzzo e del Molise “G. Caporale”, Teramo, Italy; Ospedale San Raffaele s.r.l. via olgettina; University of Oslo; Microbiology and Virology Unit, Fondazione IRCCS Policlinico San Matteo, Pavia, Italy; Institute of Medical Microbiology, University of Zurich, Zurich, Switzerland; Centro de Investigación Tibaitatá de AGROSAVIA; Nuffield Department of Medicine, University of Oxford, Oxford, UK; The Milner Centre for Evolution, Department of Life Sciences, University of Bath, BA2 7AY, UK; Department of Medical Microbiology, Stavanger University Hospital, Stavanger, Norway; Cardiff University; National Hospital for Tropical Diseases, Hanoi, Vietnam; Hanoi Medical University; Department of Pharmaceutical Microbiology, University of Ibadan; Norwegian Veterinary Institute; Statens Serum Institut, Denmark; Houston Methodist, Weill Cornell Medical College; Michael G. DeGroote Institute for Infectious Disease Research and Department of Biochemistry & Biomedical Sciences, McMaster University, Hamilton, Canada; Central Research Laboratory - Kempegowda Institute of Medical Sciences; University of Lisbon; Wellcome Sanger Institute; University Hospital of North Norway; Institut Pasteur de Bangui; European Bioinformatics Institute (EMBL-EBI); Norwegian National Advisory Unit on Detection of Antimicrobial Resistance, Department of Microbiology and Infection Control, University Hospital of North Norway, Tromsø, Norway; Department of Medical Biology, Faculty of Health Sciences, UiT The Arctic University of Norway, Tromsø, Norway; Ineos-Oxford Institute, University of Oxford; Department of Biology and Biotechnology, University of Pavia, Italy; Research Institute for Tropical Medicine, Department of Health, Manila, Philippines; Department of Pharmacy, Faculty of Health Sciences, UiT The Arctic University of Norway, Tromsø, Norway; Indiana University School of Medicine

**Keywords:** SHV, beta-lactamase, *Klebsiella pneumoniae*, extended-spectrum beta-lactamase (ESBL), beta-lactamase inhibitor resistance, prediction, antimicrobial resistance, genotype

## Abstract

Interpreting phenotypes of *bla*_SHV_ alleles in *Klebsiella pneumoniae* genomes is complex. While all strains are expected to carry a chromosomal copy conferring resistance to ampicillin, they may also carry mutations in chromosomal *bla*_SHV_ alleles or additional plasmid-borne *bla*_SHV_ alleles that have extended-spectrum β-lactamase (ESBL) activity and/or β-lactamase inhibitor (BLI) resistance activity. In addition, the role of individual mutations/amino acid changes is not completely documented or understood. This has led to confusion in the literature and in antimicrobial resistance (AMR) gene databases (e.g., NCBI’s Reference Gene Catalog and the β-lactamase database (BLDB)) over the specific functionality of individual SHV protein variants. Therefore, identification of ESBL-producing strains from *K. pneumoniae* genome data is complicated.

Here, we reviewed the experimental evidence for the expansion of SHV enzyme function associated with specific amino-acid substitutions. We then systematically assigned SHV alleles to functional classes (wildtype, ESBL, BLI-resistant) based on the presence of these mutations. This resulted in the re-classification of 37 SHV alleles compared with current assignments in NCBI’s Reference Gene Catalog and/or BLDB (21 to wildtype, 12 to ESBL, 4 to BLI-resistant). Phylogenetic and comparative genomic analyses support that; i) SHV-1 (encoded by *bla*_SHV-1_) is the ancestral chromosomal variant; ii) ESBL and BLI-resistant variants have evolved multiple times through parallel substitution mutations; iii) ESBL variants are mostly mobilised to plasmids; iv) BLI-resistant variants mostly result from mutations in chromosomal *bla*_SHV_. We used matched genome-phenotype data from the KlebNET-GSP Genotype-Phenotype Group to identify 3,999 *K. pneumoniae* isolates carrying one or more *bla*_SHV_ alleles but no other acquired β-lactamases, with which we assessed genotype-phenotype relationships for *bla*_SHV_. This collection includes human, animal, and environmental isolates collected between 2001 to 2021 from 24 countries across six continents. Our analysis supports that mutations at Ambler sites 238 and 179 confer ESBL activity, while most omega-loop substitutions do not. Our data also provide direct support for wildtype assignment of 67 protein variants, including eight that were noted in public databases as ESBL. We reclassified these eight variants as wildtype, because they lack ESBL-associated mutations, and our phenotype data support susceptibility to 3GCs (SHV-27, SHV-38, SHV-40, SHV-41, SHV-42, SHV-65, SHV-164, SHV-187).

The approach and results outlined here have been implemented in Kleborate v2.4.1 (a software tool for genotyping *K. pneumoniae* from genome assemblies), whereby known and novel *bla*_SHV_ alleles are classified based on causative mutations. Kleborate v2.4.1 was also updated to include ten novel protein variants from the KlebNET-GSP dataset and all alleles in public databases as of November 2023. This study demonstrates the power of sharing AMR phenotypes alongside genome data to improve understanding of resistance mechanisms.

**Impact statement:** Since every *K. pneumoniae* genome has an intrinsic SHV β-lactamase and may also carry additional mobile forms, the correct interpretation of *bla*_SHV_ genes detected in genome data can be challenging and can lead to *K. pneumoniae* being misclassified as ESBL-producing. Here, we use matched *K. pneumoniae* genome and drug susceptibility data contributed from dozens of studies, together with systematic literature review of experimental evidence, to improve our understanding of *bla*_SHV_ allele variation and mapping of genotype to phenotype. This study shows the value of coordinated data sharing, in this case via the KlebNET-GSP Genotype-Phenotype Group, to improve our understanding of the evolutionary history and functionality of *bla*_SHV_ genes. The results are captured in an open-source AMR dictionary utilised by the Kleborate genotyping tool, that could easily be incorporated into or used to update other tools and AMR gene databases. This work is part of the wider efforts of the KlebNET-GSP group to develop and support a unified platform tailored for the analysis and interpretation of *K. pneumoniae* genomes by a wide range of stakeholders.

**Data summary:** *Bla*_SHV_ allele sequences and class assignments are distributed with Kleborate, v2.4.1, DOI:10.5281/zenodo.10469001. **Table S1** provides a summary of *bla*_SHV_ alleles, including primary accessions, class-modifying mutations, and supporting evidence for class assignments that differ from NCBI’s Reference Gene Catalog or BLDB. Whole genome sequence data are publicly available as reads and/or assemblies, individual accessions are given in **Table S2**; corresponding genotypes and antibiotic susceptibility phenotypes and measurements are available in **Tables S3** and **S4**, respectively.

## Introduction

*Klebsiella pneumoniae* are typically resistant to ampicillin due to the production of a chromosomally-encoded Ambler class A β-lactamase enzyme, SHV. Indeed, the European Committee on Antimicrobial Susceptibility Testing (EUCAST) terms this ‘expected resistance’ (formerly ‘intrinsic resistance’) and recommends against phenotypic testing of ampicillin resistance in *K. pneumoniae*, as a susceptible result is likely to be incorrect.

While *bla*_SHV_ is a core chromosomal gene in *K. pneumoniae*, it has been mobilized out of the *K. pneumoniae* chromosome at least twice via IS*26* transposition^1^, into multiple plasmid backbones^2^ that, in turn, have spread between bacterial species. Chromosomal and plasmid forms of SHV have undergone allelic diversification to generate variants with differing functional activity, including extended-spectrum β-lactamases (ESBLs) conferring resistance to third-generation cephalosporins (3GCs) and alleles conferring resistance to β-lactamase inhibitors (BLIs).

The ESBL phenotype is facilitated through overexpression of IS*26*^3^ (e.g., SHV-2 and SHV-12). Consequently, *bla*_SHV_ genes identified in species other than *K. pneumoniae* are typically mobile and confer ESBL activity due to IS*26*, leading to a general conflation of SHV enzymes with ESBL. The existence of *bla*_SHV_ alleles with different activity profiles, including the potential for both chromosomal and plasmid-encoded genes with differing functions in a single isolate, has created confusion and can lead to an incorrect interpretation of the phenotypic impact of the molecular detection of *bla*_SHV_ genes in *K. pneumoniae*.

It is not surprising that confusion exists around the interpretation of SHV variants, given the circuitous routes through which current understanding of the origins and evolution of the enzyme have emerged. First described in 1972 as a plasmid-encoded protein of *Escherichia coli* str. 453 conferring resistance to ampicillin^4^, the enzyme was later given the name sulfhydryl-variable (SHV)-1 and was reported in multiple *Klebsiella*, *E. coli* and *Proteus mirabilis* isolates^5^. A 1979 study reported the *bla*_SHV-1_ gene as being chromosomally located in several *Klebsiella*, with a second plasmid-borne copy in one strain^6^. That study also demonstrated the transposition of the *bla*_SHV-1_ gene into different plasmid backbones, and reported the detection of *bla*_SHV-1_ on naturally-occurring plasmids of diverse types^6^. A naturally-occurring plasmid-encoded variant, designated SHV-2^7^, displayed ESBL activity and conferred resistance to 3GCs, was reported in *K. pneumoniae*, *K. ozaenae* (now known as *K. pneumoniae* subsp. *ozaenae*) and *Serratia marcescens* in 1983^8^. SHV-2 differs from SHV-1 by a single substitution (Gly to Ser) at Ambler codon 238 (amino acid 213 of the mature protein), which is sufficient to change its spectrum of activity^9^. *Bla*_SHV-2_ and *bla*_SHV-12_ are well known to be plasmid-borne, following transposition from the *K. pneumoniae* chromosome by IS*26*^1^, and found outside *K. pneumoniae*^10,11^. By 1997, twelve protein variants of SHV had been reported, most of them ESBL and mostly in *K. pneumoniae*^2^. These were designated consecutive numbers (SHV-3, −4, etc.) with the exception of the ESBL variant SHV-2a, so named due to its similar kinetic properties to SHV-2, although it is not derived from SHV-2^12^.

Besides codon 238S, amino acid substitutions identified as conferring ESBL activity mostly affect the omega-loop of SHV, including substitutions at Ambler position 179 (SHV-8^13^, SHV-24^14^) or 169 (SHV-57^15^) and an insertion at 163 (SHV-16^16^). The first variant displaying resistance to BLI (clavulanate, tazobactam) was SHV-10, which owes its unique phenotype to a substitution at codon 130^17^. Other reported BLI-resistance substitutions are located at Ambler codons 69 (SHV-49^18^), 234 (SHV-56^19^, SHV-72^20^) and 235 (SHV-107^21^).

In the sequencing era, new *bla*_SHV_ alleles are reported frequently and number in the hundreds (as of November 2023, up to *bla*_SHV-232_ have been assigned by the NCBI’s Reference Gene Catalog^22^, https://www.ncbi.nlm.nih.gov/pathogens/refgene/). Most of these alleles are also catalogued in the β-Lactamase Database^23^ (BLDB) and Comprehensive Antibiotic Resistance Database^11^ (CARD). Additions to these databases are based on novel amino acid sequences, without a requirement for biochemical characterization of enzyme function^10,24^. Database curators attempt to assign β-lactamase alleles to functional groups according to their reported spectrum of activity: narrow spectrum, extended-spectrum (ESBL), and/or BLI-resistant.

Unfortunately, the primary literature used to support functional classifications vary widely in terms of experimental design. In some cases, the presence of a *bla*_SHV_ allele in a *K. pneumoniae* isolate displaying ESBL activity has been used to ascribe ESBL functionality to the SHV enzyme, without ruling out the presence of other ESBL enzymes. This has led to the assignment of chromosomal variants with wildtype activity being reported in the literature as ESBL variants^25^ (e.g. SHV-27, SHV-41), where the error propagated to multiple AMR gene databases (in November 2023, SHV-27 and SHV-41 were still recorded as ESBL in BLDB and CARD, despite the error being reported in a 2006 publication^25^). Other examples are summarised in **Table S1** (see column ‘Evidence’ for a discussion of discrepancies between databases).

Neubauer *et al.*^26^ recently sought to clarify the role of specific substitutions in SHV functionality by systematically reconstructing isogenic mutants carrying individual substitutions (identified in naturally occurring variants with modified activity). Mutations were introduced into the *bla*_SHV-1_ background, and the spectrum of enzyme activity assessed in an *E. coli* strain lacking any other β-lactamase^26^. This confirmed the role of some substitutions in conferring ESBL activity (at Ambler site 238) or BLI resistance (at Ambler sites 69, 234, 240), however, some mutants could not be generated. In 2021, we used these findings, together with a review of experimental evidence from the literature, to systematically assign *bla*_SHV_ alleles to functional classes. We also incorporated the resulting *bla*_SHV_ database and list of functionally relevant mutations in the Kleborate tool (v2.0) for genotyping of *K. pneumoniae* genomes^27^.

Here, we aimed to systematically assess the evidence for enzyme activity by exploring genotype-phenotype relationships for naturally occurring *bla*_SHV_ alleles using a diverse set of 3,999 *K. pneumoniae* isolates with matched genome-phenotype data, which lack non-*bla*_SHV_ alleles. We also explored the evolutionary relationships and genetic context of *bla*_SHV_ alleles, with the aim of further clarifying the emergence and spread of SHV variants.

## Methods

### SHV reference database

We curated an updated set of *bla*_SHV_ nucleotide alleles for Kleborate based on a comparison of CARD^11^, NCBI’s Reference Gene Catalog^22^, and BLDB (as of November 2023, up to allele number *bla*_SHV-228_). *Bla*_SHV-6_ and *bla*_SHV-10_ were excluded as the published nucleotide sequence for *bla*_SHV-6_ is incomplete, and there is no nucleotide sequence for *bla*_SHV-10_ (only an amino acid sequence for SHV-10 which yields no exact matches to any 6-frame translations of nucleotide sequences in NCBI using tblastn). *Bla*_SHV-11_, *bla*_SHV-28_ and *bla*_SHV-31_ were represented by two nucleotide sequences each (labelled .v1, .v2) and the rest by a single nucleotide sequence. In addition to reporting matches to known alleles and the corresponding functional class, Kleborate specifically checks for and reports mutations of known functional relevance (listed in **Figure 1**; plus Ambler position 130, which was found in two novel alleles and was reported as responsible for BLI resistance in SHV-10^17^).

**Figure 1.**
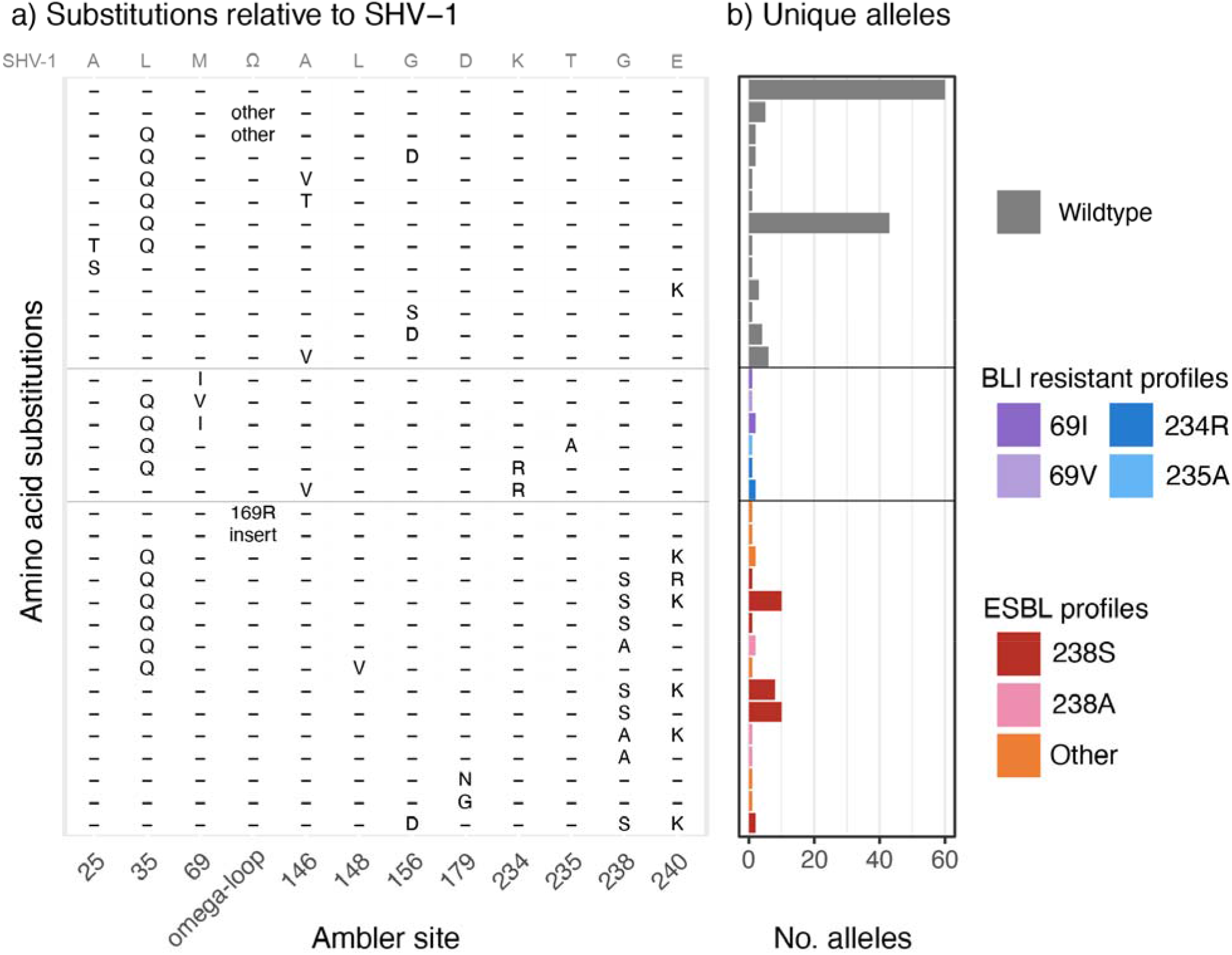
Amino acid substitution profiles associated with n=181 *bla*_SHV_ alleles. (a) Positions studied in Neubauer *et al* 2020 and tracked in Kleborate v2 are shown as columns; these include sites where substitutions have clear association with ESBL activity (148, 179, 238, 240, omega-loop (position 164-178)) or β-lactamase inhibitor (BLI) resistance activity (69, 234, 235), plus some sites (25, 35, 146, 156) that are associated with increased MIC to ceftaroline but not ceftriaxone or inhibitor resistance. Position 179 is also a part of the omega-loop and is specifically separated to show its association with ESBL profiles. Position 130 is not included, as it is found only in SHV-10 (BLI-resistant) for which there is no nucleotide sequence available. Each row indicates a unique combination of amino acids across these variable sites. The amino acids present in SHV-1 are indicated in gray at the top of the panel, where the omega-loop sequence (Ω) is RWETELNEALPGDARD. (b) The number of unique nucleotide alleles associated with each amino acid profile (row) are shown as a barplot. Colours indicate the functional class assigned to these alleles, on the basis of the mutations shown here and supporting literature (cited in the text and in **Table S1**).

We used the rules previously established for Kleborate (v2.0), as described in Supplementary Note 3 of Lam et al., 2021^27^, to assign functional classes (wildtype, ESBL, or BLI-resistant). This relied primarily on the presence of well-supported SHV substitutions as noted above (ESBL: mutation at Ambler sites 238, 179, 169; BLI resistance: mutation at sites 69, 130, 234), or direct experimental evidence for individual *bla*_SHV_ alleles (detailed in **Results**). The curated set of allele sequences and their assignment to functional classes, is included in a new release of Kleborate (v2.4.1, DOI:10.5281/zenodo.10469001) and **Table S1**. **Table S1** also includes information on each allele extracted from NCBI’s Reference Gene Catalog (including PubMed ID, Subclass, protein and nucleotide accessions), BLDB (including phenotype, sequence accessions, and alternative names) and CARD (Antibiotic Resistance Ontology [ARO] identifiers and sequence accessions).

Our curated class assignments were compared with those of BLDB and NCBI’s Reference Gene Catalog, which were interpreted as follows: wildtype (Phenotype ‘2b’ in BLDB, Subclass ‘BETA-LACTAM’ in NCBI), ESBL (Phenotype ‘2be’ in BLDB, Subclass ‘CEPHALOSPORIN’ in NCBI), and BLI-resistant (‘2br’ in BLDB, Subclass ‘BETA-LACTAM’ in NCBI but with ‘Product name’ typically including the prefix ‘inhibitor-resistant’). Discrepancies between our class assignments and those of BLDB or NCBI’s Reference Gene Catalog were reported in **Table S1**, which includes a summary of evidence based on the literature review and the phenotype data presented in this study.

Novel alleles, each encoding for unique amino acid sequences, identified in our sequence data (described below) were submitted to NCBI’s Reference Gene Catalog to obtain allele numbers in November 2023. These *bla*_SHV_ alleles (encoding SHV-233 to 237 and SHV-239 to 243), together with twelve additional *bla*_SHV_ alleles present in NCBI in November 2023 but not yet in our database (encoding SHV-115, SHV-116, SHV-132, SHV-146, SHV-171, SHV-190, SHV-191, SHV-202, and SHV-229 to 232), were also added to Kleborate v2.4.1. We also updated the database to include a single representative sequence per allele in Kleborate v2.4.1. For *bla*_SHV-11_, we selected GenBank accession AY293069 (as per BLDB, labelled .v1 in earlier versions of Kleborate), as this sequence is the correct length, and we confirmed exact matches in >1,000 of our *K. pneumoniae* genomes in >300 STs whilst .v2 was not detected. In NCBI’s Reference Gene Catalog and CARD, *bla*_SHV-11_ is represented by a different sequence (GenBank accession X98101.1) that has additional nucleotides at the start and end, differs from AY293069 at two synonymous mutations, and had no exact matches in *K. pneumoniae* whole genomes. For *bla*_SHV-28_ we used GenBank accession AF299299.1 (as per CARD and BLDB; formerly .v2 in earlier versions of Kleborate) and for *bla*_SHV-31_ we used GenBank accession AY277255.2 (as per NCBI’s Reference Gene Catalog, CARD, BLDB; formerly .v2 in Kleborate).

### Matched genome and phenotype data for *K. pneumoniae* species complex isolates

A global collection of *K. pneumoniae* species complex genomes with matched antibiotic susceptibility testing (AST) data was aggregated by the KlebNET-GSP AMR Genotype-Phenotype project group (summarised in **Tables S2-4**). This collection includes human, animal, and environmental isolates collected between 2001 to 2021 from 24 countries across six continents. Genomes were assembled from Illumina reads using Unicycler (v0.4.8) (accessions and assembly metrics in **Table S2**) and analysed using Kleborate (v2.2.0) (results in **Table S3**), which identifies the presence of acquired resistance genes/alleles including SHV, SHV protein variants, and porin defects associated with AMR^28^ (i.e., loss of OmpK35 or OmpK36, and insertions in loop 3 of OmpK36). All assemblies met the pre-agreed KlebNET-GSP criteria of <5% contamination (assessed using KmerFinder^29^ (v3.2)): Kleborate-designated species match of “strong”, ≤500 contigs; genome size 4,969,898– 6,132,846 bp; G+C content in the range 56.35%–57.98%. Contig metrics across the dataset were: contig count, mean 131.5, standard deviation (sd) 83.1; N50, mean 376272.4 bp, sd 557,147.6 bp; genome size, mean 5,418,671.9 bp, sd 163,789.2 bp; G+C content, mean 57.3%, sd 0.2%. We included only *K. pneumoniae* genomes with an exact amino acid sequence match to one or more known *bla*_SHV_ alleles, and excluded those in which other (i.e., non-SHV) β-lactamases were detected (total n=3,999 isolates for analysis).

The available antimicrobial susceptibility testing (AST) data were determined by the contributing laboratories using a range of methods, including disk diffusion, agar dilution, broth microdilution, and semi-automated methods (Vitek 2 or Phoenix), that were performed based on CLSI or EUCAST guidelines. AST data were shared in the form of disk diffusion zone sizes or minimum inhibitory concentrations (MICs). We interpreted as “susceptible” (S), “intermediate” (I), or “resistant” (R) using the EUCAST (v13.0) or Clinical and Laboratory Standards Institute (CLSI, M100 33rd edition) breakpoints, as appropriate to the assay used. As there were very few isolates categorised as I, we grouped I/R (i.e. non-wildtype) together and refer to this group as “resistant” (data in **Table S4**).

### Phylogenetic analysis

*Bla*_SHV_ nucleotide sequences were aligned using MAFFT^30^ (v6.861). Pairwise distances between aligned nucleotide sequences were calculated using the ‘dist.dna’ function in the ‘ape’ package (v5.7-1) for R (v4.2.3), and a phylogeny inferred using the BioNJ algorithm in the same package. The minimum spanning tree was inferred using GrapeTree^31^ (v1.5.0) using MSTreeV2. R packages ggplot2 (v3.4.4) and ggtree^32^ (v3.6.2) were used for data visualization.

### Genomic context of SHV alleles

Genomic location (chromosome or mobile) of *bla*_SHV_ allelic variants was determined using a combination of literature review, the CARD Prevalence, Resistomes & Variants database (v3.0.9), and BLASTN (100% nucleotide identity and coverage) searches of *bla*_SHV_ allelic variants in publicly available complete genome sequences and assembly graphs of *K. pneumoniae*. At the time of the search, CARD Prevalence, Resistomes, & Variants^11^ (v3.0.9, accessed October 2021) included 874 and 5466 *Klebsiella pneumoniae* chromosomes and plasmids, respectively, from NCBI Genomes. The presence of *bla*_SHV_ alleles amongst these chromosomes and plasmids was extracted from the CARD database. In addition, complete *K. pneumoniae* genomes (n=1296 chromosomes and n=4217 plasmids) were downloaded from NCBI using ncbi-genome-download (v0.3.1) to directly examine the genomic context of *bla*_SHV_ alleles. We used Kleborate^27^ (v2.2.0) to search for SHV variants across complete *K. pneumoniae* genomes from NCBI and the KlebNET-GSP collection.

To investigate the genetic context of mobile *bla*_SHV_ variants (represented in this analysis by *bla*_SHV-2_, *bla*_SHV-12_, *bla*_SHV-30_), we used a subset of the publicly available complete genome sequences of *K. pneumoniae* from NCBI. We selected 30 random genomes from different STs (determined by Kleborate (v.2.2.0), and ran Mauve^33^ (v2015-02-25) to identify the collinear block containing *bla*_SHV_. We then used BLASTN to search for this collinear block in all publicly available complete chromosome sequences of *K. pneumoniae* from NCBI, to confirm its broader conservation. To visualise the genetic context of mobile variants of *bla*_SHV_ compared with the typical chromosomal context from which they were presumably mobilised, we extracted 10 kbp of sequence upstream and downstream of the gene from genomes CP103302.1 (*bla*_SHV-2_), NC_009650.1 (*bla*_SHV-12_) NZ_CP017936.1 (*bla*_SHV-30_) and NZ_CP032170.1 (*bla*_SHV-30_). We then used Prokka^34^ (v1.14.6) and clinker^35^ (v0.0.24) to annotate and visually compare the extracted genomic regions with a representative sequence of the chromosomal collinear block extracted from the chromosome of *K. pneumoniae* strain MGH 78578 (accession CP000647.1). In addition, we used flankophile^36^ (v0.2.10) to extract *bla*_SHV_ flanking regions (5 kbp upstream and downstream) across the previously used NCBI’s publicly available complete chromosome *K. pneumoniae* sequences to capture genetic variation in *bla*_SHV_ flanking regions. We then used CD-HIT-EST^37^ (v4.8.1) to cluster the flanking regions with l790% nucleotide sequence similarity. We used the same visualisation methods as described above. We also investigated the presence of insertion elements 10 kbp upstream of wildtype *bla*_SHV_ in the genomes of phenotypically 3GC resistant isolates using the BLASTN and the ISfinder^38^ database.

For the set of complete genomes, the *bla*_SHV_ copy number was calculated based on the number of unique non-overlapping BLASTN hits. For the matched genotype-phenotype dataset, the copy number of *bla*_SHV_ in draft genomes was estimated by analysing Illumina read sets, calculating the ratio of read depth for *bla*_SHV_ vs the mean read depth of the seven *K. pneumoniae* loci used for multi-locus sequence typing, using SRST2^39^ (v0.2.0) to perform the mapping and depth calculations. Copy number estimates are included along with other genotype information in **Table S3**.

## Results

### Distribution of activity-modifying mutations

In Kleborate v2.2.0’s database, there are a total of 181 unique *bla*_SHV_ alleles, corresponding to 178 unique protein sequences or variants (**Table S1**). **Figure 1** illustrates the distribution of key amino-acid substitutions (hereafter the term mutations is used for both nucleotide and amino acid variation for convenience, even though amino acid changes are a consequence of the actual mutations) across these alleles. Among the *bla*_SHV_ alleles identified, 38 encoded mutations relative to SHV-1 at Ambler positions 238 (n=36) and 179 (n=2). These specific mutations have been observed by Neubauer *et al.*^26^ to confer 3GC resistance, classifying these variants as ESBL. Five additional protein variants were assigned as ESBL based on primary literature reports (**Table S1**): SHV-16^16^ (omega-loop insertion between Ambler sites 167-168), SHV-57^15^ (omega-loop substitution 169R), SHV-31 (encoded by divergent alleles SHV-31.v1 and SHV-31.v2^40^, each carrying mutations 35Q and 240K), SHV-70^41^ (mutation 148V). Eight alleles harboured a substitution at a site associated with BLI-resistance^26^ and were classified accordingly: Ambler site 69 (SHV-49^18^, SHV-52, SHV-92, SHV-203), or 234 (SHV-56^19^, SHV-72^20^, SHV-73). SHV-107 (harbouring mutation 235A) was also classified as BLI-resistant based on primary literature^21^. The remaining 130 alleles were assigned as wildtype.

### Evolutionary relationships and genomic context

To understand the evolutionary relationships between *bla*_SHV_ alleles, we inferred a cladogram and minimum spanning tree from the nucleotide sequence alignment (**Figure 2, Figure S1**). Pairwise genetic distances between allele sequences support *bla*_SHV-1_ as the ancestral form, as it has the smallest distance to all other variants (mean 5.1 substitutions, total distance 918; compared with mean 5.4, total 969 for *bla*_SHV-11_ which had the next lowest values). We therefore rooted the phylogeny at *bla*_SHV-1_. We identified a chromosomal *bla*_SHV_ collinear block (7,585 bp) conserved in 90.5% (n=1,236/1,366) of complete *K. pneumoniae* genomes with >90% nucleotide identity and >90% coverage (**Figure 3**). In general, there is low genetic variation of the chromosomal *bla*_SHV_ flanking regions (5 kbp upstream and downstream), where 95.1% (n=1,229/1,292) of complete genomes had l790% nucleotide sequence similarity in their flanking regions. Comparing the chromosomal *bla*_SHV_ collinear block with the genomic context of plasmid-borne and IS*26*-mediated *bla*_SHV-2_ and *bla*_SHV-12_, the chromosomal *bla*_SHV_ collinear block is conserved in the genomic context of *bla*_SHV-2_ and flanked by IS*26*. Similarly, the chromosomal *bla*_SHV_ collinear block is partially conserved (59% coverage, 99% identity of 7,585 bp) in the *bla*_SHV-12_ genomic context. The gene directly downstream of *bla*_SHV_, *glpR*, encodes a glycerol-3-phosphate regulon repressor, which is conserved across the genomic contexts of chromosomal and plasmid-borne *bla*_SHVs._

**Figure 2.**
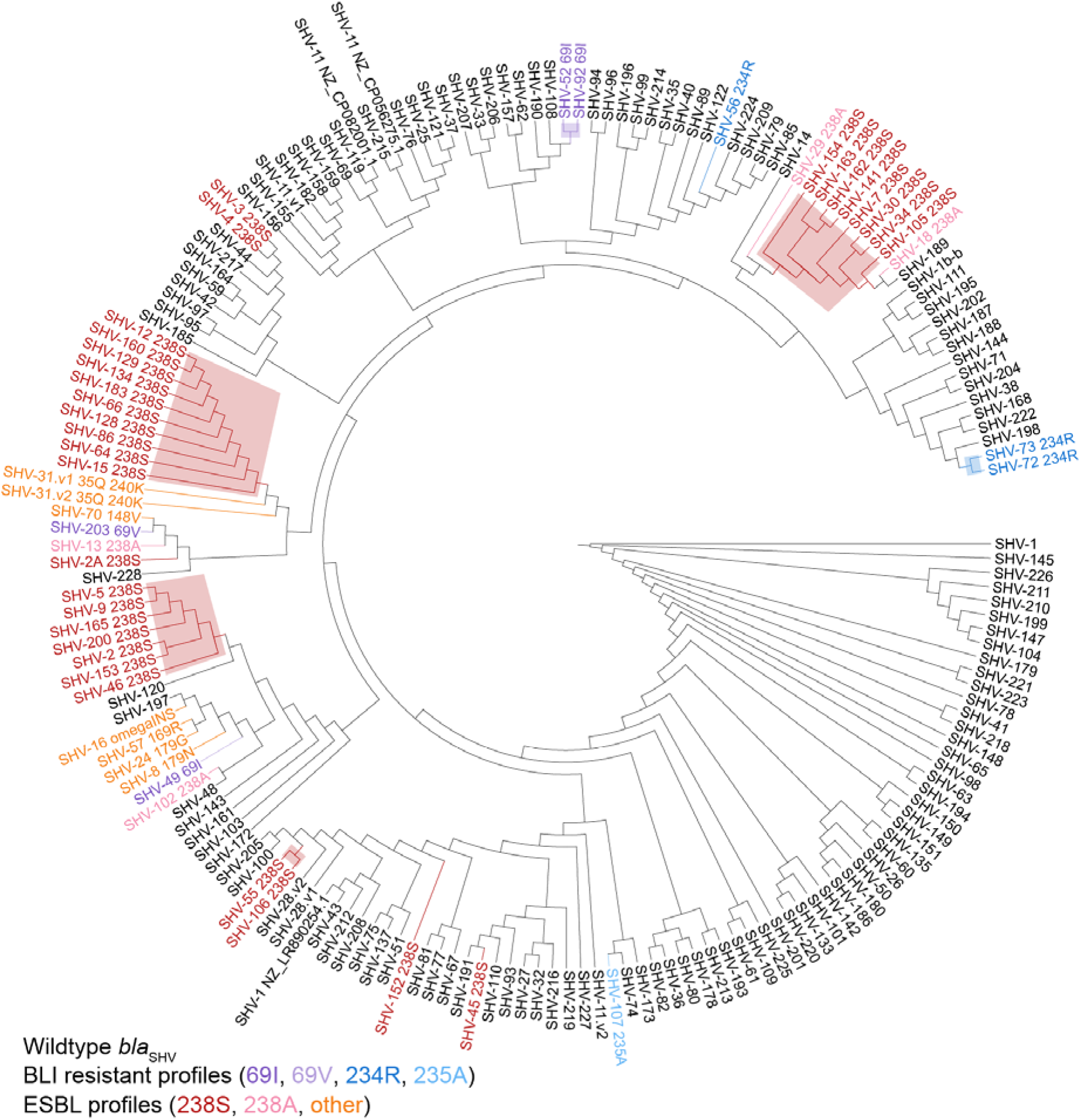
Cladogram for n=181 *bla*_SHV_ alleles. The cladogram was inferred from a pairwise genetic distance matrix calculated from nucleotide sequences using BioNJ, rooted on SHV-1. Tips are labelled with the SHV allele name, and coloured to indicate the mutation profile (black=wildtype; red, orange, pink = ESBL profiles; blue, purple = BLI resistant profiles). For alleles classed as non-wildtype, the class-modifying mutation is included in the label (e.g. 238S indicates substitution of serine at Ambler site 238 in the encoded protein; ‘omegaINS’ refers to a 6-amino acid insertion in the omega-loop between Ambler codons 167-168). Shading indicates clusters of alleles referred to in the text, which may share class-modifying mutations via vertical inheritance. SHV-1 NZ_LR890254.1, SHV-11 NZ_CP056275.1, SHV-11 NZ_CP082001.1 are nucleotide variants, but have the same protein sequences as SHV-1 and SHV-11, respectively.

**Figure 3.**
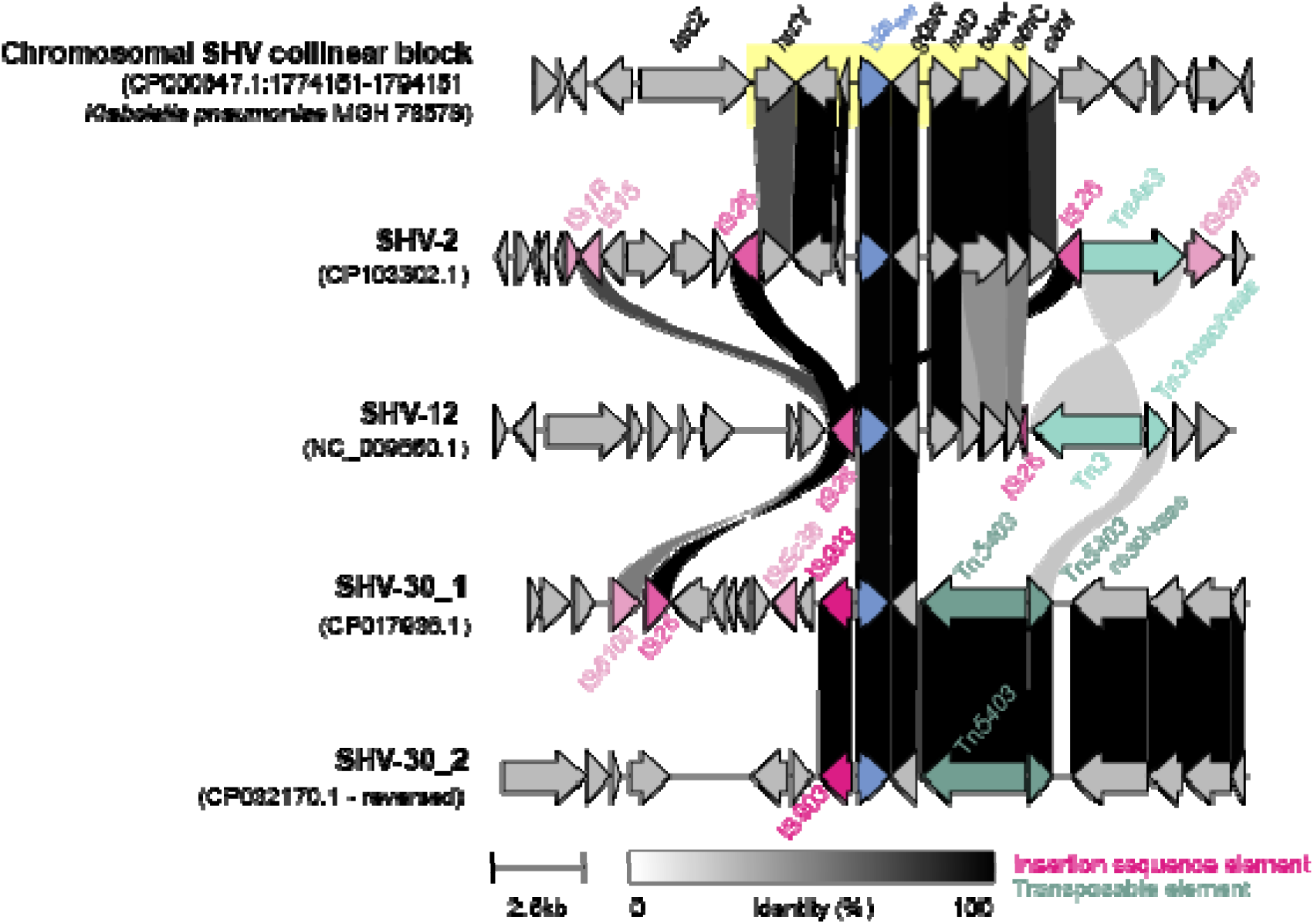
A comparison of the genomic context of SHV allele clusters. Upstream (10kbp) and downstream (10kbp) sequences of each *bla*_SHV_ were extracted and aligned. The 7,585 bp chromosomal SHV collinear block is highlighted in yellow. *Bla*_SHV_ is coloured in blue, while mobile genetic elements, such as insertion sequences and transposons, are illustrated in pink and green, respectively. Percent identity between the genes are shown by the gradient scale bar.

ESBL alleles were distributed throughout the cladogram (pink, red, orange in **Figure 2**), consistent with at least 19 independent mutation events (n=12 in Ambler codon 238, n=1 in codon 148, n=2 in codon 179, n=2 others in the omega-loop, and n=2 [SHV-31.v1 and SHV-31.v2] in codon 240)^40^. Some ESBL alleles formed clusters that appear to share a resistance-conferring mutation (238S) via inheritance from a common ancestor (shading, **Figure 2**). These include two pairs of alleles (*bla*_SHV-3_/*bla*_SHV-4_, *bla*_SHV-55_/*bla*_SHV-106_) and three larger clusters centred around *bla*_SHV-2_/*bla*_SHV-5_ (n=7), *bla*_SHV-12_ (n=10), and *bla*_SHV-7_/*bla*_SHV-30_ (n=8). Mobilisation of *bla*_SHV-2_ and *bla*_SHV-12_ by IS*26* are well-documented^1,10^. *Bla*_SHV-3_ and *bla*_SHV-4_ are also known to be plasmid-borne^42^ and found in species outside *Klebsiella*^10^, although we could not identify a complete plasmid sequence in which to explore the specific genetic context of the mobilised region. Members of the *bla*_SHV-7_ cluster, including *bla*_SHV-7_^43^, *bla*_SHV-_^44^ and *bla*SHV-34 ^45^, have been reported as plasmid-borne and found in *Enterobacter*. Among the *bla*_SHV-7_ cluster, only *bla*_SHV-30_ was detected amongst complete *K. pneumoniae* genome sequences in NCBI. This allele was identified in two similar plasmid sequences (99.99% identity over 55,821 bp of shared sequence [85% coverage], detected in ST2938 [accession NZ_CP032170.1] and ST45 [accession NZ_CP017936.1]), where it was flanked by IS*903* and Tn*5403* (**Figure 3**). This provides a potentially novel, non-IS*26*-mediated, mobility mechanism for this ESBL cluster, which was found in a total of 13 *K. pneumoniae* genomes belonging to 7 STs (and one *K. variicola* genome) in our genome collection. We could find no evidence to support that *bla*_SHV-55_/*bla*_SHV-106_ have been mobilised out of the *K. pneumoniae* chromosome. NCBI BLAST did not identify these alleles outside *K. pneumoniae*, nor in any complete *K. pneumoniae* genomes. We identified a single instance in our genome collection (*bla*_SHV-106_ in a ST14 genome); read analysis indicated a copy number of one, suggesting this was the only copy of *bla*_SHV_ in the genome, and assembly graph analysis supported its location in the chromosome. Four of five ESBL allele clusters therefore appear to be plasmid-borne, and likely reflect diversification of ESBL alleles following mobilisation from the *K. pneumoniae* chromosome.

BLI-resistant alleles (n=8) were distributed throughout the cladogram (blue, purple in **Figure 2**), consistent with at least six independent mutation events (n=3 in Ambler codon 69, n=2 in 234, and n=1 in 235). These alleles have not been reported outside of *K. pneumoniae* and sequence searches of NCBI and CARD did not detect evidence of them in non-*K. pneumoniae* genomes. The original reports of *bla*_SHV-56_ and *bla*_SHV-49_ confirmed these variants as chromosomally located^19^, and we also found *bla*_SHV-52_ and *bla*_SHV-56_ in draft genome sequences where assembly graph inspection confirmed they were located on chromosomal contigs. The other alleles *bla*_SHV-72_, *bla*_SHV-73_ and *bla*_SHV-92_ were not found in our genome collection or in NCBI genomes via BLASTN search. The original report of *bla*_SHV-92_ states that it was detected in a transconjugant, suggesting that it was plasmid-borne^46^, however we found no evidence of any other BLI-resistant alleles being mobile. These data suggest that the currently reported BLI-resistant *bla*_SHV_ alleles have arisen in wildtype chromosomal *bla*_SHV_ backgrounds. With the exception of *bla*_SHV-92_, these BLI-resistant alleles have not yet been mobilised to plasmids, which is consistent with the low prevalence of the phenotype reported in *K. pneumoniae* isolates.

### Genotype-phenotype relationships

We compared *bla*_SHV_ alleles with AST phenotypes for 3GCs and BLIs in a set of n=3,999 *K. pneumoniae* genomes that carried at least one *bla*_SHV_ allele and no other β-lactamase (**Table S2-4**). Within these genomes, we identified 70 of the known 181 *bla*_SHV_ alleles (38% of those in the Kleborate (v2.2.0)).

Eight known ESBL protein variants (classified as such in the literature and here) were identified in isolates that were tested for susceptibility to ceftazidime and all but one (sole representative of SHV-106) showed evidence for resistance (see **Table 1**, **Figure 4**). All of these protein variants have at least a 238S substitution, with the exception of SHV-31.v1 which had both 35Q and 240K substitutions. All isolates representing the remaining protein variants and for which data were available, also showed evidence for resistance to ceftriaxone. However, resistance to cefotaxime was more variable. Eleven other isolates carried *bla*_SHV_ alleles with a non-synonymous mutation in the omega-loop, but were not previously reported as ESBL alleles (*bla*_SHV-51_, and four novel alleles, see **Table 1**); all tested susceptible to ceftazidime (**Table 1**, **Figure 4**) and all other 3GCs for which they were tested (**Table 1**).

**Figure 4.**
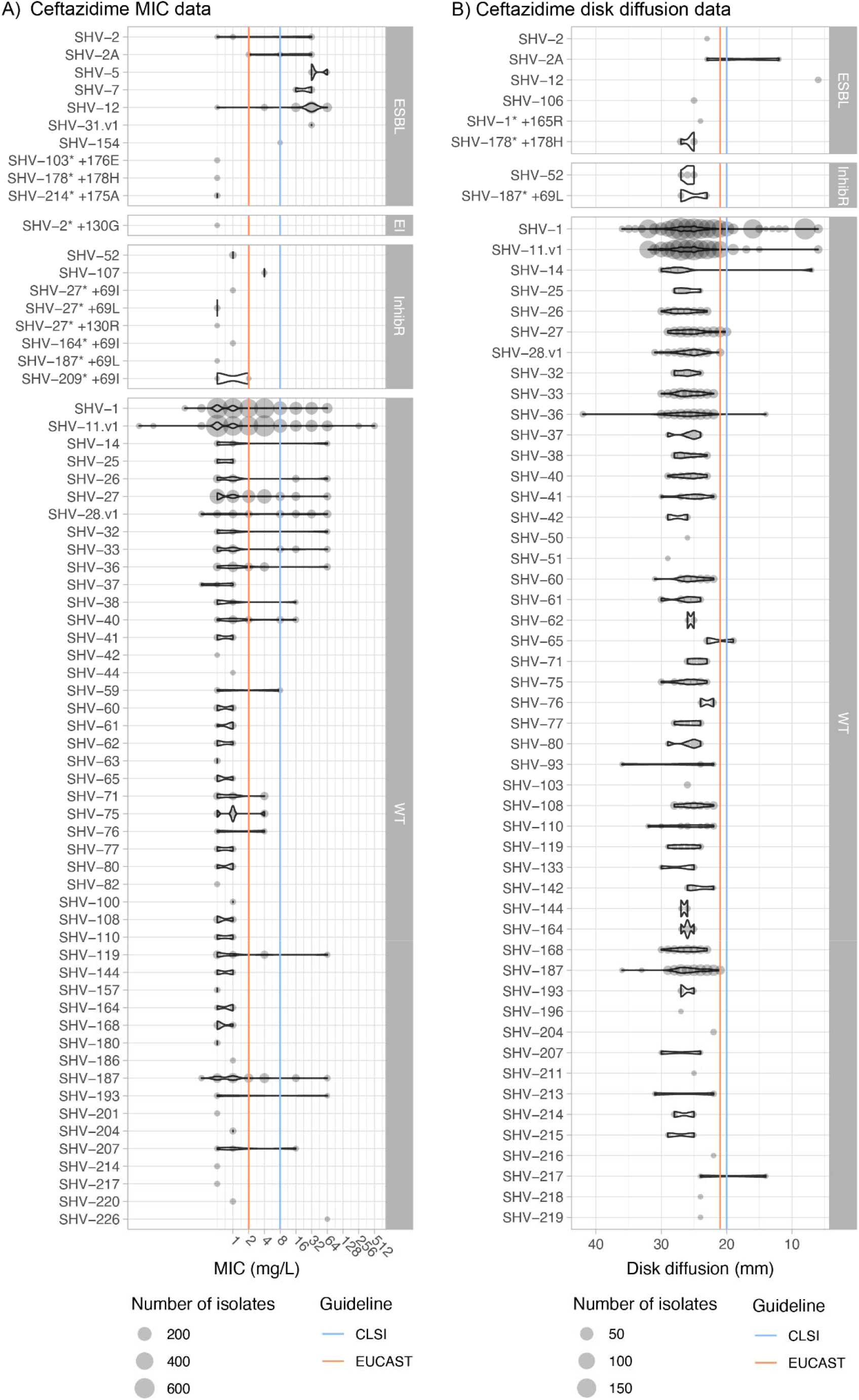
AST values distributions for ceftazidime. The size of each circle represents the number of genomes with an SHV allele and no other acquired β-lactamase. Minimum inhibitory concentration (a) and disk diffusion (b) measurements show the distribution of phenotypes for each SHV-allele. SHV alleles are grouped based on extended spectrum β-lactamase (ESBL), ESBL and β-lactamase inhibitor resistant (EI), β-lactamase inhibitor resistant (inhibR), and wildtype (WT) phenotype classifications. EUCAST (v13.0) or CLSI (M100 33rd edition) intermediate breakpoints are indicated using orange and blue lines, respectively. SHV-187* +69L is SHV-132 in Kleborate v2.4.1. For MIC values, larger values indicate increased resistance; for disk diffusion results, larger zone sizes indicate increased susceptibility.

**Table 1.**
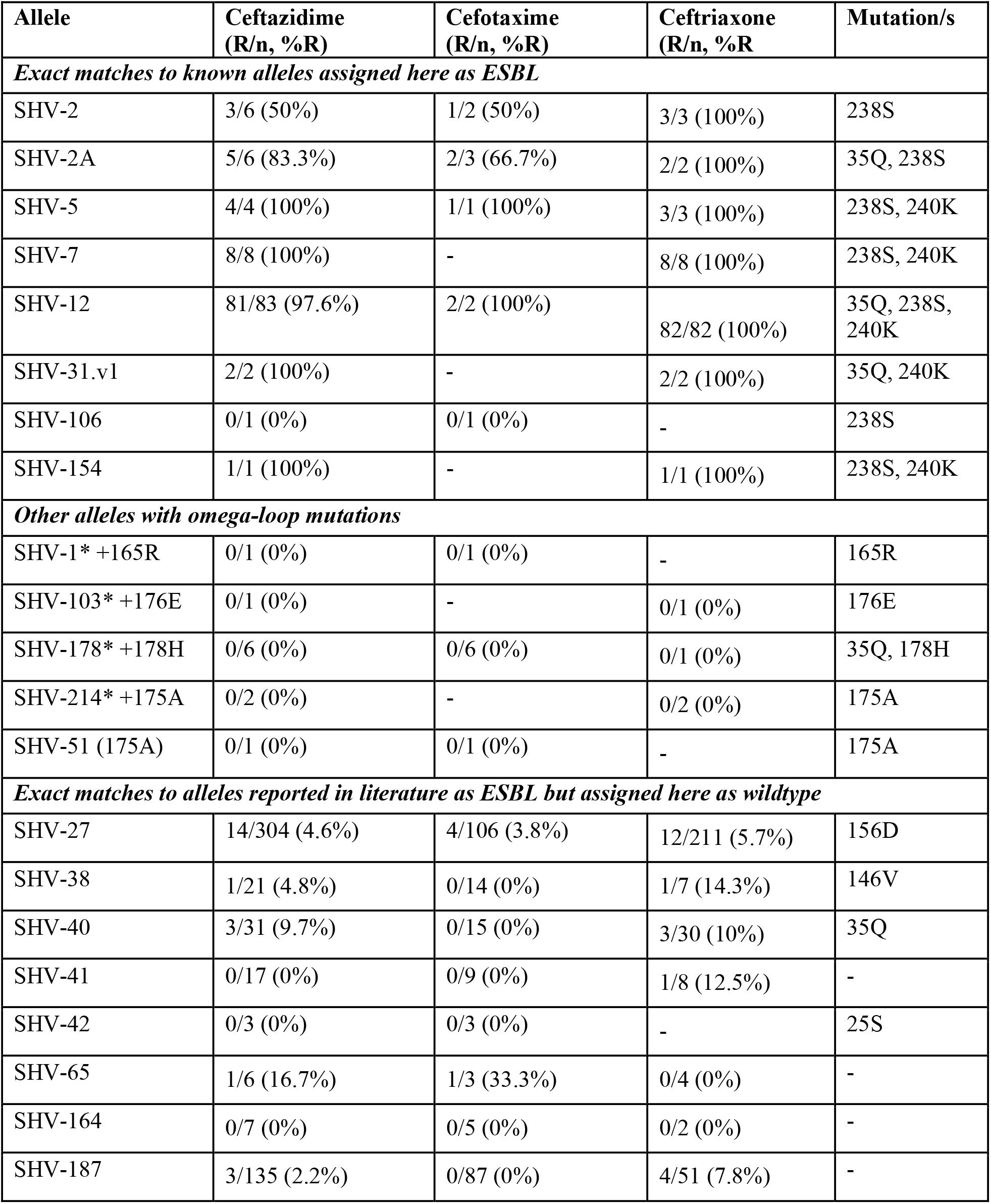
Third-generation cephalosporin susceptibility phenotypes for ESBL-assigned alleles. Novel alleles identified in this study have been highlighted with an ‘*’.

We identified n=533 isolates with alleles that were initially reported as ESBL and assigned as such in NCBI’s Reference Gene Catalog and/or BLDB, but do not carry any causative mutations and were therefore classified in our database as wildtype (encodes SHV-27, SHV-38, SHV-40, SHV-41, SHV-42, SHV-65, SHV-164, SHV-187). Our phenotype data support the assignment to wildtype for all these alleles (see **Table 1**). A summary of the comparison with BLDB and NCBI’s Reference Gene Catalog’s class assignments is given in **Table S1**.

Two BLI-resistant variants were identified in isolates that were tested for susceptibility to piperacillin-tazobactam and/or amoxicillin-clavulanic acid: SHV-52 (which harbours 69I) and SHV-107 (which harbours 235A) (see **Table 2**). The two isolates carrying SHV-107 came from the same study and were resistant to amoxicillin-clavulanic acid as expected (MIC 32 mg/L via the automated Vitek platform; piperacillin-tazobactam results were not available). All isolates carrying SHV-52 and tested for piperacillin-tazobactam were susceptible (n=12, from five different studies using either disk diffusion or MIC via Vitek); n=9 of these isolates were also tested for amoxicillin-clavulanic acid, all were susceptible.

**Table 2.** Beta-lactamase inhibitor susceptibility phenotypes for inhibitor-resistance-assigned alleles. Novel alleles identified in this study have been highlighted with an ‘*’. ^†^SHV-187* +69L is SHV-132 in Kleborate v2.4.1.

| Allele | Piperacillin-tazobactam (R/n, %R) | Amoxicillin-clavulanic acid (R/n, %R) | Mutation/s |
| --- | --- | --- | --- |
| ***Exact matches to known alleles assigned as inhibitor-resistant*** |  |  |  |
| SHV-52 | 0/12 (100%) | 0/9 (100%) | 35Q, 69I |
| SHV-107 | - | 2/2 (100%) | 235A |
| ***Other alleles with mutations at Ambler site 69, 130, 234 or 235*** |  |  |  |
| SHV-2* +130G | 1/1 (100%) | 1/1 (100%) | 130G, 238S |
| SHV-27* +69I | 0/1 (0%) | 0/1 (0%) | 69I |
| SHV-27* +69L | 0/2 (0%) | 0/2 (0%) | 69L |
| SHV-27* +130R | 0/1 (0%) | 0/1 (0%) | 130R |
| SHV-110* +69I | 1/1 (100%) | 1/1 (100%) | 35Q, 69I, 156D |
| SHV-164* +69I | 0/1 (0%) | 0/1 (0%) | 69I |
| SHV-187* +69L<sup>†</sup> | 0/2 (0%) | 0/1 (0%) | 69L |
| SHV-209* +69I | 2/2 (100%) | 1/1 (100%) | 35Q, 69I |
| ***Exact matches to alleles reported in literature as inhibitor-resistant but assigned here as wildtype*** |  |  |  |
| SHV-26 | 13/63 (20.6%) | 10/59 (16.9%) | - |

The closely-related allele SHV-92, which shares the 69I mutation and clusters with *bla*_SHV-52_ in the cladogram (differing from it at a single nucleotide, see **Figure 1**), was not present in our dataset so we could not assess its phenotype directly; the original report of this allele also did not assess phenotype^46^. Nine isolates carried novel variants harbouring a substitution at Ambler site 69; six of these tested susceptible to BLIs and three tested resistant to piperacillin-tazobactam and amoxicillin-clavulanic acid (**Table 2**, **Figure 5**). The resistant isolates were: n=1 carrying a novel variant closest to SHV-110 with additional mutation 69I (accession: SRR15097887), and n=2 (from different studies^47,48^, accessions: SRR15098057, ERR486441) harbouring a novel variant closest to SHV-209 with additional mutation 69I (**Table 2, Table S4**). Two isolates were identified with novel alleles carrying mutations at codon 130, one (carrying SHV-27 plus 130R) was tested for susceptibility to BLIs but showed susceptibility to both piperacillin-tazobactam and amoxicillin-clavulanic acid via disk diffusion (**Table 2**, **Figure 5**). The other isolate (carrying SHV-2 plus 130G) was resistant to both piperacillin-tazobactam and amoxicillin-clavulanic acid via agar dilution (**Table 2**, **Figure 5**).

**Figure 5.**
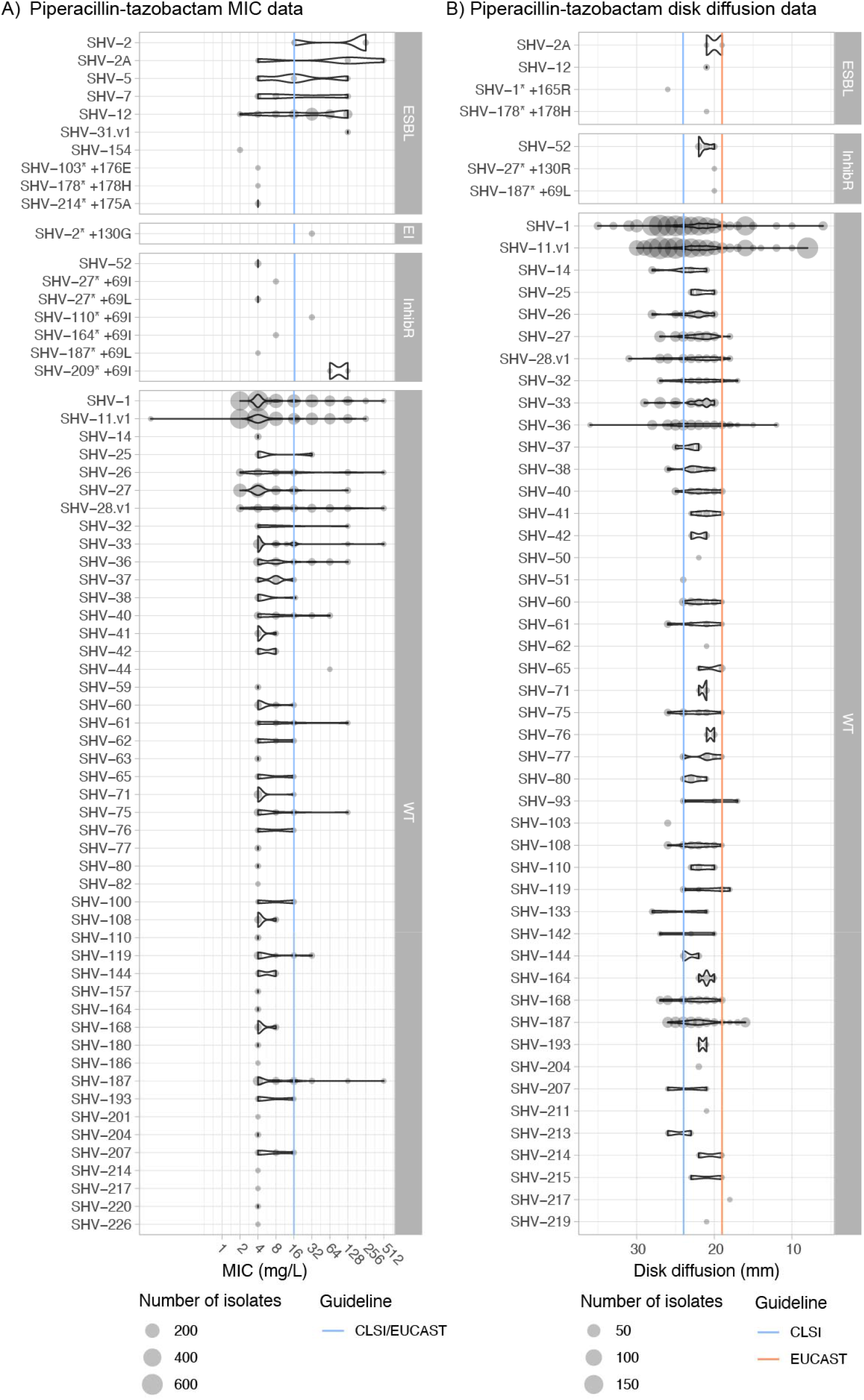
AST values distributions for piperacillin-tazobactam. The size of each circle represents the number of genomes with an SHV allele and no other acquired β-lactamase. Minimum inhibitory concentration (a) and disk diffusion (b) measurements show the distribution of phenotypes for each SHV-allele. SHV alleles are grouped based on extended spectrum β-lactamase (ESBL), extended spectrum β-lactamase and β-lactamase inhibitor resistant (EI), β-lactamase inhibitor resistant (inhibR), and wildtype (WT) phenotype classifications. EUCAST (v13.0) or CLSI (M100 33rd edition) intermediate breakpoints are indicated using orange and blue lines, respectively. SHV-187* +69L is SHV-132 in Kleborate v2.4.1. For MIC values, larger values indicate increased resistance; for disk diffusion results, larger zone sizes indicate increased susceptibility.

We also identified n=63 isolates carrying SHV-26, which we assigned as wildtype due to lacking functional mutations, but is classified as BLI resistant (2br) in BLDB. The original report of SHV-26^49^ described it as harbouring a mutation at Ambler site 187 and reduced susceptibility to amoxicillin-clavulanic acid (to ‘intermediate’ levels). However this mutation (A187T) was tested by Neubauer *et al.*^26^, who found no effect on BLI susceptibility and concluded the phenotype was likely incorrectly assigned. Our data show some evidence of a BLI-resistant phenotype (n=13/63 (20.6%) to piperacillin-tazobactam, n=10/59 (17%) resistant to amoxicillin-clavulanic acid), but with majority support for wildtype (**Table 2**).

Sixty alleles classified as wildtype were detected in the genome collection (total n=3858 isolates), and the wildtype phenotype was supported in all cases. Thirty-six of these alleles (60%) were found only in 3GC-susceptible isolates. One allele (*bla*_SHV-59_) was found in one resistant isolate and one susceptible (both ST76 with no other resistance determinants detected). The remaining n=23 wildtype-classified alleles were primarily found in susceptible strains (66.7–98.0% susceptible, per allele). These include alleles *bla*_SHV-1_ and *bla*_SHV-11_, the most common and well-known wildtype alleles. We hypothesised that increased copy number of *bla*_SHV_ and/or porin mutations could explain 3GC and BLI resistance in isolates with wildtype-assigned *bla*_SHV_ alleles and no other acquired β-lactamases. Amongst isolates with a wildtype-assigned *bla*_SHV_ allele, *bla*_SHV_ copy number was indeed significantly associated with ceftazidime MIC (correlation = 0.21, p<1×10^-15^ using linear regression on log_2_ MIC), disk diffusion zone diameter (correlation = −0.76, p<1×10^-15^ using linear regression), and clinical resistance (mean 2.7 vs 1.1 copies, p=2×10^-6^ using Wilcoxon rank sum test). Similarly, *bla*_SHV_ copy number in isolates with wildtype *bla*_SHVs_ was significantly associated with piperacillin-tazobactam MIC (correlation = 0.25, p<1×10^-15^ using linear regression on log_2_ MIC), disk diffusion zone diameter (correlation = −0.10, p<1×10^-15^ using linear regression), and clinical resistance (mean 2.3 vs 1.1 copies, p=2×10^-6^ using Wilcoxon rank sum test). The presence of two or more copies of *bla*_SHVs_ was significantly associated with ceftazidime resistance (OR 3.6, p=6×10^-13^ amongst isolates with a wildtype-assigned *bla*_SHV_ allele), accounting for 25.6% of the resistance observed amongst these isolates. A further 12.8% of ceftazidime resistance could potentially be explained by porin defects in isolates with a single *bla*_SHV_ copy (see **Figure 6**). We also investigated the presence of insertion sequences upstream of wildtype *bla*_SHV_ (with no other acquired β-lactamases) in genomes of phenotypically 3GC resistant isolates that could potentially explain the phenotype^3^, but there were none identified. For piperacillin-tazobactam, presence of two or more copies of *bla*_SHV_ was significantly associated with resistance (OR 4.78, p<1×10^-15^ amongst isolates with a wildtype-assigned *bla*_SHV_ allele), accounting for 34.1% of unexplained resistance, with a further 9.5% potentially explained by porin defects (see **Figure 7**).

**Figure 6.**
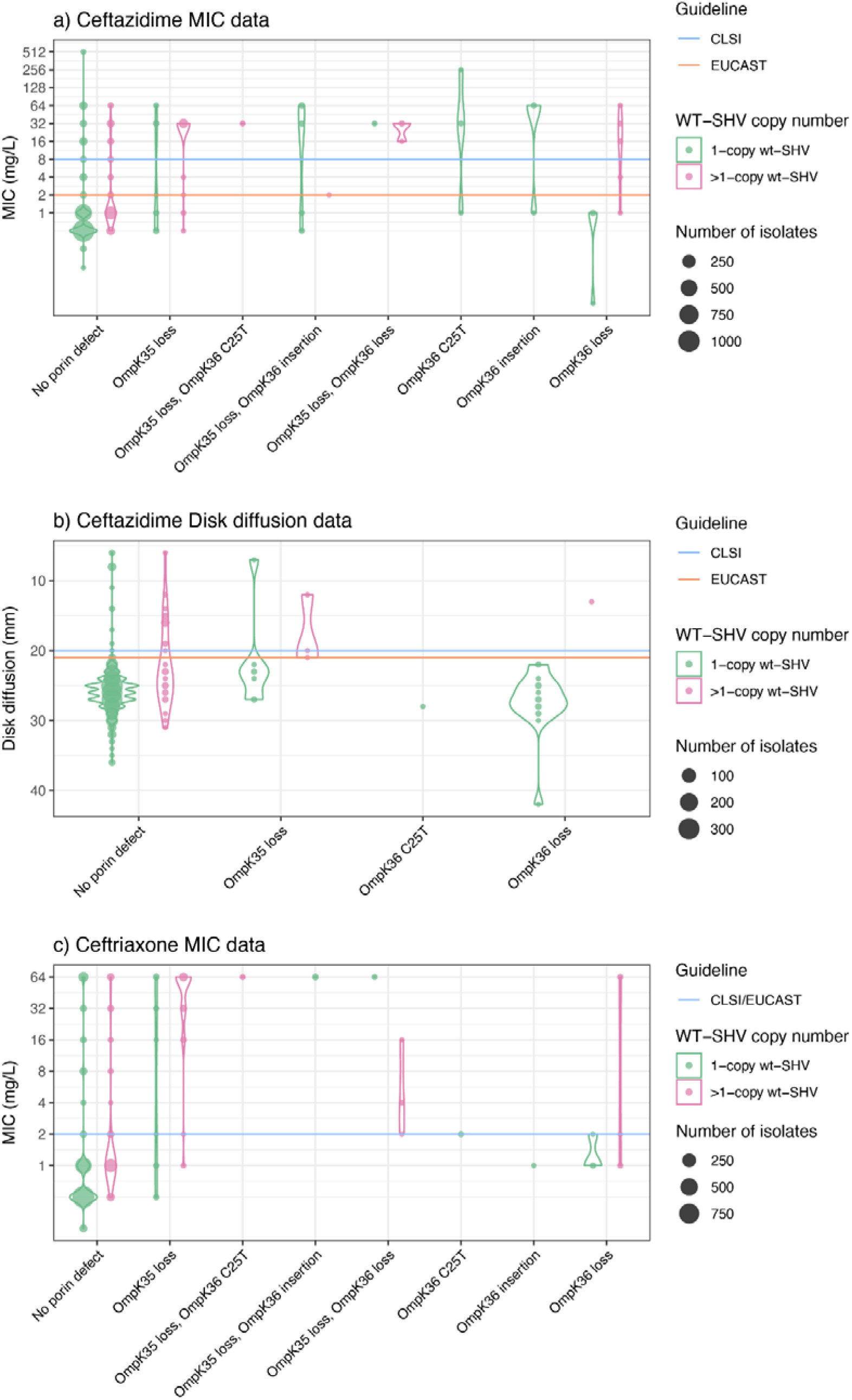
Presence of porin defects and copy number effects amongst isolates with wildtype-assigned alleles with genomes tested against 3GCs. Barplots show the distribution of susceptibility testing measures, coloured by copy number and porin defects that are tracked by Kleborate (as per inset legend), for wildtype SHV alleles (n=1659 isolates tested against ceftazidime and n=1937 isolates tested against ceftriaxone). EUCAST (v13.0) or CLSI (M100 33rd edition) intermediate/resistant breakpoints are indicated using orange and blue lines, respectively.

**Figure 7.**
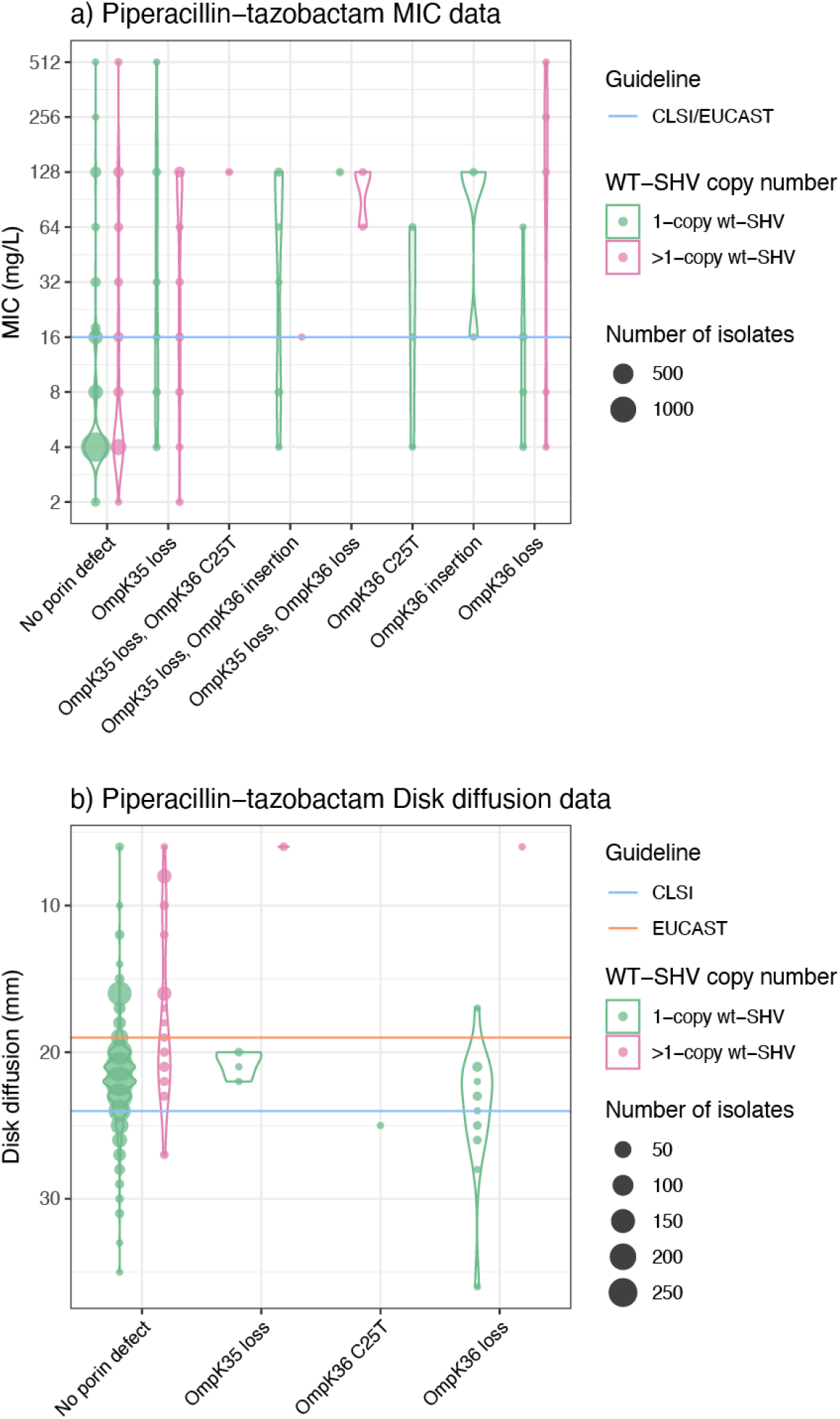
Presence of porin defects and copy number effects amongst isolates with wildtype-assigned alleles with genomes tested against piperacillin-tazobactam. Barplots show the distribution of susceptibility testing measures, coloured by porin defects that are tracked by Kleborate (as per inset legend), for n=2268 isolates with wildtype SHV alleles. EUCAST (v13.0) or CLSI (M100 33rd edition) intermediate/resistant breakpoints are indicated using orange and blue lines, respectively.

## Discussion

*Bla*_SHV_ alleles have been studied since their discovery in 1972 and were first explored phylogenetically in 1990 to study the context of *bla*_SHV-2_ and its relationships with other β-lactamase genes^50^. As new *bla*_SHV_ variants are discovered, phylogenetic trees were inferred to explore their ancestry and relationships with each other^2,10,51^. Most recently, Liakopoulos *et al* inferred a maximum likelihood tree with 149 SHV-type β-lactamases, but it was unclear which *bla*_SHV_ was the likely ancestral variant^10^. It has been assumed that SHV-1 is the ancestral variant since it was the first *bla*_SHV_ discovered and our cladogram, pairwise distance data and minimum-spanning tree also support *bla*_SHV-1_ as the ancestral variant (**Figure 2, Figure S1**). There is also support from Chaves *et al.*^52^ and Haeggman *et al.*^53^, who show that *bla*_SHV-1_ is predominantly species-specific to *K. pneumoniae* and has a long evolutionary history as a stable chromosomal gene, suggesting that even the ancestor of *bla*_SHV-1_ is also from the *K. pneumoniae* chromosome.

Our phylogenetic and comparative genomic analyses support that ESBL and BLI-resistant variants of *bla*_SHV_ have evolved multiple times independently through parallel substitution mutations (**Figure 2**), and that many of these variants have been mobilized out of the *K. pneumoniae* chromosome via independent events (**Figure 3**), enabling them to spread between lineages, species, and genera. We found evidence of mobilisation for most ESBL variants, but only one BLI-resistance conferring variant (SHV-92). Consistent with this, most 3GC-resistant *K. pneumoniae* carrying ESBL variants and no other β-lactamases were found to have multiple copies of SHV (presumably a chromosomal copy with wildtype activity plus a plasmid-borne copy with ESBL activity).

We have reviewed the classification of *bla*_SHV_ alleles into functional classes to better support the interpretation of genomic data. Our work builds on the experimental study of Neubauer *et al*., which provided evidence of the role of specific mutations to enzyme activity. By systematically assigning alleles to functional classes based on the presence of specific mutations associated with enzyme activity (**Figure 1**), rather than presence in an ESBL or BLI-resistant isolate (which may confuse mobile and chromosomal variants), we propose re-classification of 20 *bla*_SHV_ alleles from ESBL to wildtype (n=12 changes vs NCBI’s Reference Gene Catalog, n=14 changes vs BLDB, see **Table S1**).

We used matched genotype-phenotype data, for 3,999 *K. pneumoniae* carrying *bla*_SHV_ and no other acquired β-lactamases, to assess predictability of phenotype based on *bla*_SHV_ alleles (**Figures 4-5**, **Tables 1-2**). For this we used our Kleborate tool to identify and type *bla*_SHV_ alleles, and specific SHV mutations associated with a change in enzyme activity. This analysis provided additional support for the role of 238S and 179G^13,14^ in ESBL activity and consequent 3GC resistance, but suggests that most changes in the omega-loop do not result in a change in activity. These data also support our classification of variants SHV-27, SHV-38, SHV-40, SHV-41, SHV-42, SHV-65, SHV-164, and SHV-187 – which lack mutations at site 238 or any other mutations associated experimentally with resistance – as wildtype.

Mutations 69I and 69V have been thought to explain BLI resistance of variants SHV-49, SHV-52, SHV-92 and SHV-203, respectively, and were found by Neubauer *et al* to confer resistance to piperacillin-tazobactam. Interestingly, our data do not support a simple association between Ambler site 69 mutations and BLI resistance in *K. pneumoniae*, whether in the SHV-52 variant (n=0/12 resistant to piperacillin-tazobactam or amoxicillin-clavulanic acid) or arising *de novo* in other SHV backgrounds (SHV-27, SHV-110, SHV-164, SHV-187, SHV-209) (n=3/9 resistant). We identified two genomes with a mutation at Ambler site 235 (both SHV-107, which carry mutation 235A) which were both resistant to amoxicillin-clavulanic acid (piperacillin-tazobactam was not tested), providing support for the role of this mutation, which was confirmed by Neubauer *et al*^26^.

The approach and results outlined here have been implemented in Kleborate v2.4.1, along with all new alleles identified in this study, and all those available in public databases as of 7 November 2023. In the Kleborate v2.4.1 database, known *bla*_SHV_ alleles classified as ESBL are those with amino acid substitutions at Ambler site 238 (n=36 alleles), 179 (SHV-8), 169 (SHV-57), 148 (SHV-70), 240K+35Q (SHV-31) or insertion in the omega-loop (SHV-16). SHV variants are classified as BLI-resistant if they possess mutations at Ambler site 69, 130, 234 or 235. Where exact nucleotide or protein matches are found to a known allele, these are reported in the relevant column (Bla_ESBL, Bla_inhibR, Bla_ESBL_inhibR, Bla_wt) based on the classification in the Kleborate database. As the mutations noted above are considered causative of a change of enzyme activity (class-modifying), Kleborate checks for these mutations in all SHV sequences and reports them in a separate column, SHV_mutations. If a class-modifying mutation is detected in an otherwise wildtype-classified allele background, the novel allele will be reported in the relevant functional column, i.e., Bla_ESBL, Bla_ESBL_inhibR, or Bla_inhibR rather than Bla_wt, and labelled with the mutation.

Kleborate will also report any mutation in the omega-loop (sites 164-179) in the SHV_mutations column, as it is theoretically possible that any modification disrupting the omega-loop structure could impact function^16,54,55^. However, detection of these mutations will not change the class assignment in Kleborate since most changes are likely to be non-functional and all novel omega-loop mutants we identified in our study tested susceptible to 3GCs (**Table 2**). Our phenotype data also do not support a simple association between mutations at Ambler site 69 and clinical resistance to piperacillin-tazobactam or amoxicillin-clavulanic acid (**Table 2**). However, our numbers are small (n=9 isolates, of which three tested resistant), and the functional evidence for BLI resistance associated with mutations at this site are convincing^18,26,56^, therefore we consider it appropriate to distinguish alleles with Ambler site 69 mutations from wildtype alleles in the Kleborate database and reporting.

The KlebNET-GSP matched genotype-phenotype dataset yielded coverage of 40% of known *bla*_SHV_ alleles in otherwise β-lactamase-free backgrounds, which is essential to interpret the role of *bla*_SHV_ specifically. Despite other alleles being in unfavourable genomic contexts, our approach enabled a systematic assessment of how *bla*_SHV_ alleles are assigned to functional classes in the public AMR gene databases and provides evidence that some existing assignments are incorrect (**Table S1**). In turn this helped us to implement a more transparent and consistent approach to detecting and reporting known and novel *bla*_SHV_ alleles in *K. pneumoniae* genomes, via Kleborate v2.4.1. In addition, the diversity of this dataset (isolates from 24 different countries across 2001-2021 and collected from humans, animals, and environments) avoids the very often local epidemiological effects that could bias results. Additional insights into the role of genetic background, expression, and co-expression of SHV variants and/or other β-lactamases on resistance mechanisms will help to further clarify the impact of individual variants and lead to better interpretation of genotypes and prediction of phenotypes.

This study exemplifies the importance of sharing AST data together with genome data, and the potential role for global collaboration such as KlebNET-GSP to utilise this data to enhance understanding of resistance mechanisms. This is particularly relevant in cases like *bla*_SHV_, where complex evolutionary processes have contributed to the emergence and mobilisation of resistant variants within and between the originating species. As the KlebNET-GSP isolate collection grows, we intend to regularly update this analysis to support the growing evidence for *bla*_SHV_ phenotypes; to explore genotype-phenotype variation in the homologous enzymes of other members of the *K. pneumoniae* species complex (*bla*_OKP_ in *K. quasipneumoniae* and *bla*_LEN_ in *K. variicola*), and to undertake similar analyses to inform understanding of the mechanisms of resistance to other drug classes relevant to treatment of *K. pneumoniae* infection.

## Acknowledgements

KlebNET-GSP AMR Genotype-Phenotype Group (https://klebnet.org/) shares this work on behalf of the Kp-T7 and Kp-MDR studies, The Norwegian Study Group on *Klebsiella pneumoniae*, Kp-NORM study, JPI-AMR consortium SpARK, NIHR Global Health Research Unit Genomic Surveillance of Antimicrobial Resistance, the ‘Controlling Superbugs’ flagship study and the Victorian Carbapenemase-Producing Enterobacterales (CPE) program, Vietnam ICU WGS study, GHRU-GSA-Ibadan, and REDUCEAMU project.

## Authors contributions

Conceptualization – K.E.H.; Methodology – M.M.C.L., K.K.T., K.E.H.; Software – M.M.C.L., K.E.H.; Validation – M.M.C.L., K.K.T., K.E.H., Formal analysis – K.K.T.; Investigation – K.K.T., Resources – K.E.H., M.B., S.B., K.B., S.B., A.C., D.M.C., J.C., M.C., A.C., A.C., N.D., P.D., A.E., R.F., E.J.F., A.F., C.L.G., Y.G., B.H., M.A.K.H., L.N.M.H., L.T.H., B.H., O.I., A.W.J.J., H.K., F.K., T.L., I.H.L., S.W.L., G.L., M.L., A.J.M., A.G.M., G.N., A.O.O., I.N.O., H.P., J.P., M.H.P., F.P., N.R., A.R., K.L.R.K., L.R., C.R., Ø.S., K.S., D.S., H.S., V.S., N.L.S., S.S., A.S., N.S., M.S., A.S., P.N.T., N.T., H.A.T., E.T., V.D.T., N.V.T., J.V., T.W., B.W., H.W., G.D.W., K.L.W.; Data curation – M.M.C.L., K.K.T.; Project administration – K.E.H.; Supervision – K.E.H.; Writing - Original Draft – K.K.T.; Writing - Review & Editing – K.K.T., M.M.C.L., R.R.W., K.L.W., M.B., S.B., K.B., S.B., A.C., D.M.C., J.C., M.C., A.C., A.C., N.D., P.D., A.E., R.F., E.J.F., A.F., C.L.G., Y.G., B.H., M.A.K.H., L.N.M.H., L.T.H., B.H., O.I., A.W.J.J., H.K., F.K., T.L., I.H.L., S.W.L., G.L., M.L., A.J.M., A.G.M., G.N., A.O.O., I.N.O., H.P., J.P., M.H.P., F.P., N.R., A.R., K.L.R.K., L.R., C.R., Ø.S., K.S., D.S., H.S., V.S., N.L.S., S.S., A.S., N.S., M.S., A.S., P.N.T., N.T., H.A.T., E.T., V.D.T., N.V.T., J.V., T.W., B.W., H.W., G.D.W.,K.E.H.; Visualization – K.K.T.; Funding acquisition – K.E.H.

---

**Figure S1:**
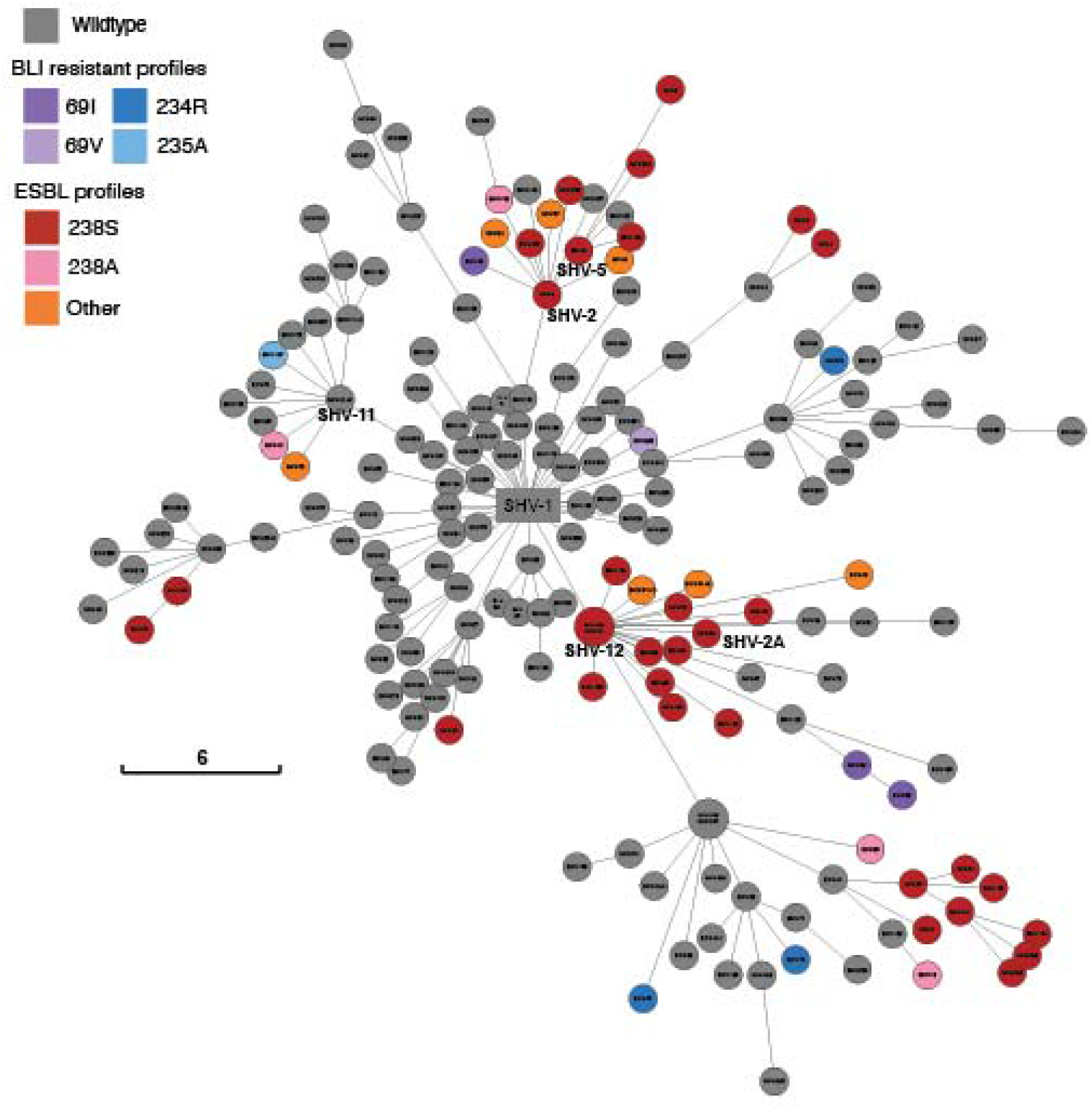
Minimum spanning tree of *bla*_SHV_ alleles. Minimum spanning tree was inferred using GrapeTree. Size of circles indicates the number of alleles at the same position. Tips are labelled with the SHV allele name, and coloured to indicate the mutation profile (grey=wildtype; red, orange, pink = ESBL mutation profiles; blue, purple = BLI resistant mutation profiles). Scale bar indicates genetic distance.

**Figure S2:**
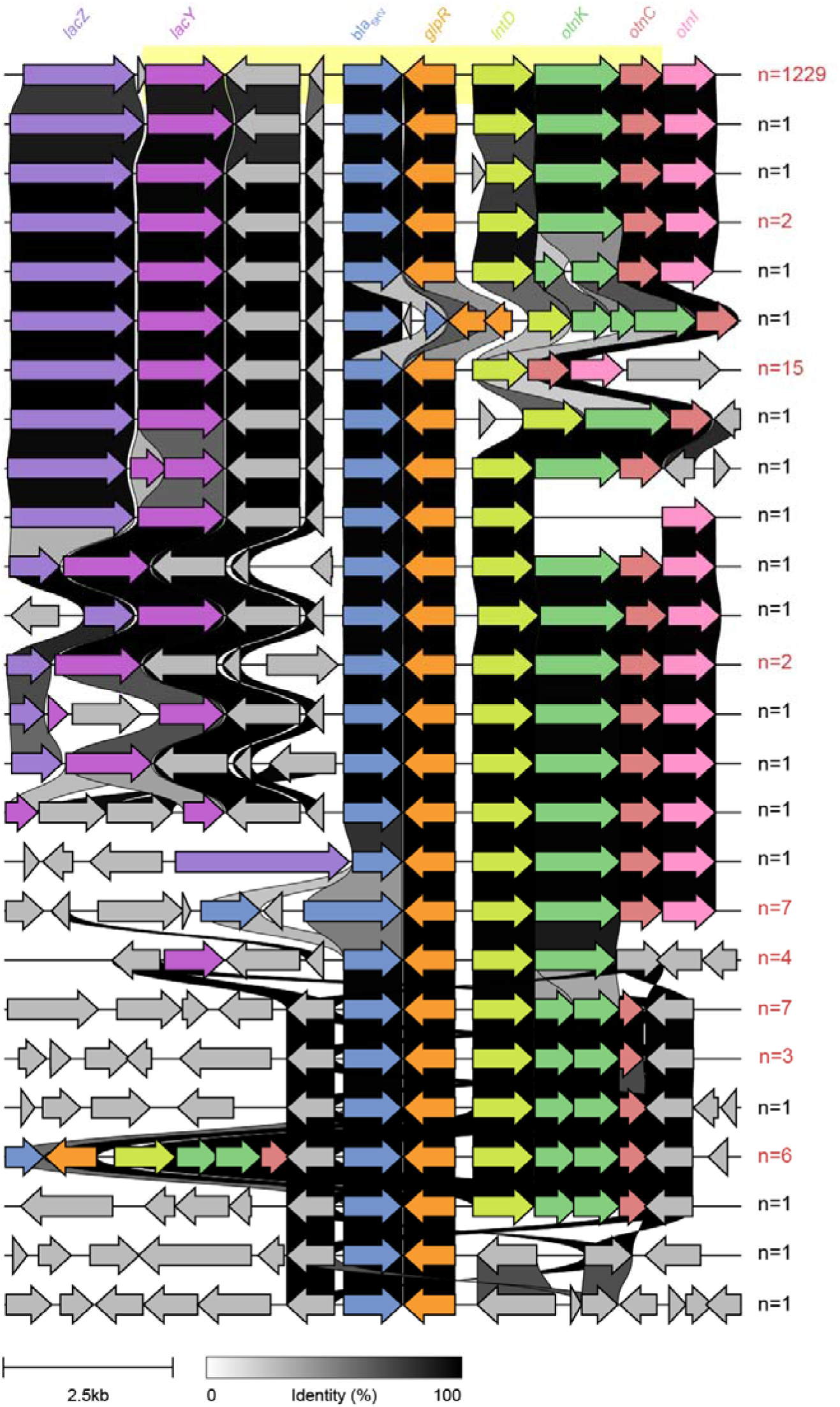
Genetic variation of *bla*_SHV_ flanking regions. Upstream (5 kbp) and downstream (5 kbp) *bla*_SHV_ were extracted and aligned. The prevalence of each flanking region (≥90% nucleotide sequence similarity) is labelled at the end of each region, where red text indicates flanking regions identified in more than one genome. The 7,585 bp chromosomal SHV collinear block is highlighted in yellow. *Bla*_SHV_ is coloured in blue, while other genes are coloured as labelled. Percent identity between the genes are shown by the gradient scale bar.

## List of supp tables and figures

**Table S1:** Curated database of SHV alleles, including primary accessions, class-modifying mutations, genomic context.

**Table S2:** Genome data, including NCBI accessions for reads or assemblies, source / metadata, assembly quality metrics.

**Table S3:** Genotypes inferred from genome data.

**Table S4:** Antimicrobial susceptibility phenotypes data.

**Figure S1:** Minimum spanning tree of *bla*_SHV_ alleles.

**Figure S2:** Genetic variation of *bla*_SHV_ flanking regions.

## References

1. Ford, P. J. & Avison, M. B. Evolutionary mapping of the SHV β-lactamase and evidence for two separate IS26-dependent blaSHV mobilization events from the Klebsiella pneumoniae chromosome. J. Antimicrob. Chemother. 54, 69–75 (2004).

2. Heritage, J., M’Zali, F. H., Gascoyne-Binzi, D. & Hawkey, P. M. Evolution and spread of SHV extended-spectrum β-lactamases inGram-negative bacteria. J. Antimicrob. Chemother. 44, 309–318 (1999).

3. Hammond, D. S., Schooneveldt, J. M., Nimmo, G. R., Huygens, F. & Giffard, P. M. bla(SHV) Genes in Klebsiella pneumoniae: different allele distributions are associated with different promoters within individual isolates. Antimicrob. Agents Chemother. 49, 256–263 (2005).

4. Pitton, J.-S. Mechanisms of bacterial resistance to antibiotics. Ergebnisse der Physiol. Rev. Physiol. Vol. 65 15–93 (1972). doi:10.1007/3-540-05814-1_2

5. Matthew, M., Hedges, R. W. & Smith, J. T. Types of beta-lactamase determined by plasmids in gram-negative bacteria. J. Bacteriol. 138, 657–662 (1979).

6. Nugent, M. E. & Hedges, R. W. The nature of the genetic determinant for the SHV-1 β-lactamase. MGG Mol. Gen. Genet. 175, 239–243 (1979).

7. Kliebe, C., Nies, B. A., Meyer, J. F., Tolxdorff-Neutzling, R. M. & Wiedemann, B. Evolution of plasmid-coded resistance to broad-spectrum cephalosporins. Antimicrob. Agents Chemother. 28, 302–307 (1985).

8. Knothe, H., Shah, P., Krcmery, V., Antal, M. & Mitsuhashi, S. Transferable resistance to cefotaxime, cefoxitin, cefamandole and cefuroxime in clinical isolates of Klebsiella pneumoniae and Serratia marcescens. Infection 11, 315–317 (1983).

9. Barthélémy, M., Péduzzi, J., Ben Yaghlane, H. & Labia, R. Single amino acid substitution between SHV-1 β-lactamase and cefotaxime-hydrolyzing SHV-2 enzyme. FEBS Lett. 231, 217–220 (1988).

10. Liakopoulos, A., Mevius, D. & Ceccarelli, D. A review of SHV extended-spectrum β-lactamases: Neglected yet ubiquitous. Front. Microbiol. 7, 219996 (2016).

11. Alcock, B. P. et al. CARD 2023: expanded curation, support for machine learning, and resistome prediction at the Comprehensive Antibiotic Resistance Database. Nucleic Acids Res. 51, D690–D699 (2023).

12. Podbielski, A., Schönling, J. A., Melzer, B. & Warnatz, K. Molecular cloning and nucleotide sequence of a new plasmid-coded Klebsiella pneumoniae beta-lactamase gene (SHV-2a) responsible for high-level cefotaxime resistance. Zentralbl. Bakteriol. 275, 369–373 (1991).

13. Rasheed, J. K. et al. Evolution of extended-spectrum beta-lactam resistance (SHV-8) in a strain of Escherichia coli during multiple episodes of bacteremia. Antimicrob. Agents Chemother. 41, 647–653 (1997).

14. Kurokawa, H. et al. A new SHV-derived extended-spectrum beta-lactamase (SHV-24) that hydrolyzes ceftazidime through a single-amino-acid substitution (D179G) in the - loop. Antimicrob. Agents Chemother. 44, 1725–1727 (2000).

15. Ma, L. et al. Novel SHV-derived extended-spectrum beta-lactamase, SHV-57, that confers resistance to ceftazidime but not cefazolin. Antimicrob. Agents Chemother. 49, 600–605 (2005).

16. Arpin, C. et al. SHV-16, a beta-lactamase with a pentapeptide duplication in the omega loop. Antimicrob. Agents Chemother. 45, 2480–2485 (2001).

17. Prinarakis, E. E., Miriagou, V., Tzelepi, E., Gazouli, M. & Tzouvelekis, L. S. Emergence of an inhibitor-resistant beta-lactamase (SHV-10) derived from an SHV-5 variant. Antimicrob. Agents Chemother. 41, 838–840 (1997).

18. Dubois, V. et al. SHV-49, a novel inhibitor-resistant β-lactamase in a clinical isolate of Klebsiella pneumoniae. Antimicrob. Agents Chemother. 48, 4466–4469 (2004).

19. Dubois, V. et al. Molecular and Biochemical Characterization of SHV-56, a Novel Inhibitor-Resistant β-Lactamase from Klebsiella pneumoniae. Antimicrob. Agents Chemother. 52, 3792–3794 (2008).

20. Mendonça, N. et al. The Lys234Arg substitution in the enzyme SHV-72 is a determinant for resistance to clavulanic acid inhibition. Antimicrob. Agents Chemother. 52, 1806–1811 (2008).

21. Manageiro, V. et al. Characterization of the inhibitor-resistant SHV β-lactamase SHV-107 in a clinical Klebsiella pneumoniae strain coproducing GES-7 enzyme. Antimicrob. Agents Chemother. 56, 1042–1046 (2012).

22. Feldgarden, M. et al. AMRFinderPlus and the Reference Gene Catalog facilitate examination of the genomic links among antimicrobial resistance, stress response, and virulence. Sci. Rep. 11, 12728 (2021).

23. Naas, T. et al. Beta-lactamase database (BLDB) – structure and function. J. Enzyme Inhib. Med. Chem. 32, 917–919 (2017).

24. Bradford, P. A. et al. Consensus on β-Lactamase Nomenclature. Antimicrob. Agents Chemother. 66, (2022).

25. Lin, T. L. et al. Extended-spectrum beta-lactamase genes of Klebsiella pneumoniae strains in Taiwan: recharacterization of shv-27, shv-41, and tem-116. Microb. Drug Resist. 12, 12–15 (2006).

26. Neubauer, S. et al. A Genotype-Phenotype Correlation Study of SHV β-Lactamases Offers New Insight into SHV Resistance Profiles. Antimicrob. Agents Chemother. 64, (2020).

27. Lam, M. M. C. et al. A genomic surveillance framework and genotyping tool for Klebsiella pneumoniae and its related species complex. Nat. Commun. 2021 121 12, 1–16 (2021).

28. Fajardo-Lubián, A., Ben Zakour, N. L., Agyekum, A., Qi, Q. & Iredell, J. R. Host adaptation and convergent evolution increases antibiotic resistance without loss of virulence in a major human pathogen. PLOS Pathog. 15, e1007218 (2019).

29. Hasman, H. et al. Rapid whole-genome sequencing for detection and characterization of microorganisms directly from clinical samples. J. Clin. Microbiol. 52, 139–146 (2014).

30. Katoh, K., Misawa, K., Kuma, K. I. & Miyata, T. MAFFT: a novel method for rapid multiple sequence alignment based on fast Fourier transform. Nucleic Acids Res. 30, 3059–3066 (2002).

31. Zhou, Z. et al. GrapeTree: visualization of core genomic relationships among 100,000 bacterial pathogens. Genome Res. 28, 1395–1404 (2018).

32. Yu, G. Using ggtree to Visualize Data on Tree-Like Structures. Curr. Protoc. Bioinforma. 69, (2020).

33. Darling, A. C. E., Mau, B., Blattner, F. R. & Perna, N. T. Mauve: Multiple Alignment of Conserved Genomic Sequence With Rearrangements. Genome Res. 14, 1394 (2004).

34. Seemann, T. Prokka: rapid prokaryotic genome annotation. Bioinformatics 30, 2068– 2069 (2014).

35. Gilchrist, C. L. M. & Chooi, Y. H. clinker & clustermap.js: automatic generation of gene cluster comparison figures. Bioinformatics 37, 2473–2475 (2021).

36. Thorn, A. V., Aarestrup, F. M. & Munk, P. Flankophile: a bioinformatic pipeline for prokaryotic genomic synteny analysis. Microbiol. Spectr. 12, (2024).

37. Fu, L., Niu, B., Zhu, Z., Wu, S. & Li, W. CD-HIT: accelerated for clustering the next-generation sequencing data. Bioinformatics 28, 3150–3152 (2012).

38. Siguier, P., Perochon, J., Lestrade, L., Mahillon, J. & Chandler, M. ISfinder: the reference centre for bacterial insertion sequences. Nucleic Acids Res. 34, (2006).

39. Inouye, M. et al. SRST2: Rapid genomic surveillance for public health and hospital microbiology labs. Genome Med. 6, 1–16 (2014).

40. Mazzariol, A., Roelofsen, E., Koncan, R., Voss, A. & Cornaglia, G. Detection of a new SHV-type extended-spectrum β-lactamase, SHV-31, in a Klebsiella pneumoniae strain causing a large nosocomial outbreak in the Netherlands. Antimicrob. Agents Chemother. 51, 1082–1084 (2007).

41. Ling, B. D. et al. Characterisation of a novel extended-spectrum beta-lactamase, SHV-70, from a clinical isolate of Enterobacter cloacae in China. Int. J. Antimicrob. Agents 27, 355–356 (2006).

42. Nicolas, M. H., Jarlier, V., Honore, N., Philippon, A. & Cole, S. T. Molecular characterization of the gene encoding SHV-3 beta-lactamase responsible for transferable cefotaxime resistance in clinical isolates of Klebsiella pneumoniae. Antimicrob. Agents Chemother. 33, 2096–2100 (1989).

43. Levison, M. E. et al. Regional occurrence of plasmid-mediated SHV-7, an extended-spectrum beta-lactamase, in Enterobacter cloacae in Philadelphia Teaching Hospitals. Clin. Infect. Dis. 35, 1551–1554 (2002).

44. Szabó, D. et al. Molecular analysis of the simultaneous production of two SHV-type extended-spectrum beta-lactamases in a clinical isolate of Enterobacter cloacae by using single-nucleotide polymorphism genotyping. Antimicrob. Agents Chemother. 49, 4716–4720 (2005).

45. Heritage, J., Chambers, P. A., Tyndall, C. & Buescher, E. S. SHV-34: an extended-spectrum beta-lactamase encoded by an epidemic plasmid. J. Antimicrob. Chemother. 52, 1015–1017 (2003).

46. Lavilla, S. et al. Prevalence of qnr genes among extended-spectrum β-lactamase-producing enterobacterial isolates in Barcelona, Spain. J. Antimicrob. Chemother. 61, 291–295 (2008).

47. Moradigaravand, D., Martin, V., Peacock, S. J. & Parkhill, J. Evolution and Epidemiology of Multidrug-Resistant Klebsiella pneumoniae in the United Kingdom and Ireland. MBio 8, (2017).

48. Sherry, N. L. et al. Genomics for Molecular Epidemiology and Detecting Transmission of Carbapenemase-Producing Enterobacterales in Victoria, Australia, 2012 to 2016. J. Clin. Microbiol. 57, (2019).

49. Chang, F. Y., Siu, L. K., Fung, C. P., Huang, M. H. & Ho, M. Diversity of SHV and TEM beta-lactamases in Klebsiella pneumoniae: gene evolution in Northern Taiwan and two novel beta-lactamases, SHV-25 and SHV-26. Antimicrob. Agents Chemother. 45, 2407–2413 (2001).

50. Huletsky, A., Couture, F. & Levesque, R. C. Nucleotide sequence and phylogeny of SHV-2 beta-lactamase. Antimicrob. Agents Chemother. 34, 1725–1732 (1990).

51. Newire, E. A., Ahmed, S. F., House, B., Valiente, E. & Pimentel, G. Detection of new SHV-12, SHV-5 and SHV-2a variants of extended spectrum Beta-lactamase in Klebsiella pneumoniae in Egypt. Ann. Clin. Microbiol. Antimicrob. 12, 16 (2013).

52. Chaves, J. et al. SHV-1 β-lactamase is mainly a chromosomally encoded species-specific enzyme in Klebsiella pneumoniae. Antimicrob. Agents Chemother. 45, 2856– 2861 (2001).

53. Hæggman, S., Löfdahl, S., Paauw, A., Verhoef, J. & Brisse, S. Diversity and evolution of the class A chromosomal beta-lactamase gene in Klebsiella pneumoniae. Antimicrob. Agents Chemother. 48, 2400–2408 (2004).

54. Kuzin, A. P. et al. Structure of the SHV-1 beta-lactamase. Biochemistry 38, 5720–5727 (1999).

55. Sampson, J. M. et al. Ligand-Dependent Disorder of the Ω Loop Observed in Extended-Spectrum SHV-Type β-Lactamase. Antimicrob. Agents Chemother. 55, 2303 (2011).

56. Giakkoupi, P. et al. Properties of Mutant SHV-5 β-Lactamases Constructed by Substitution of Isoleucine or Valine for Methionine at Position 69. Antimicrob. Agents Chemother. 42, 1281 (1998).

