## Supplementary figures and images for "Diversity, functional classification and genotyping of SHV β-lactamases in *Klebsiella pneumoniae*"

### FigureS1

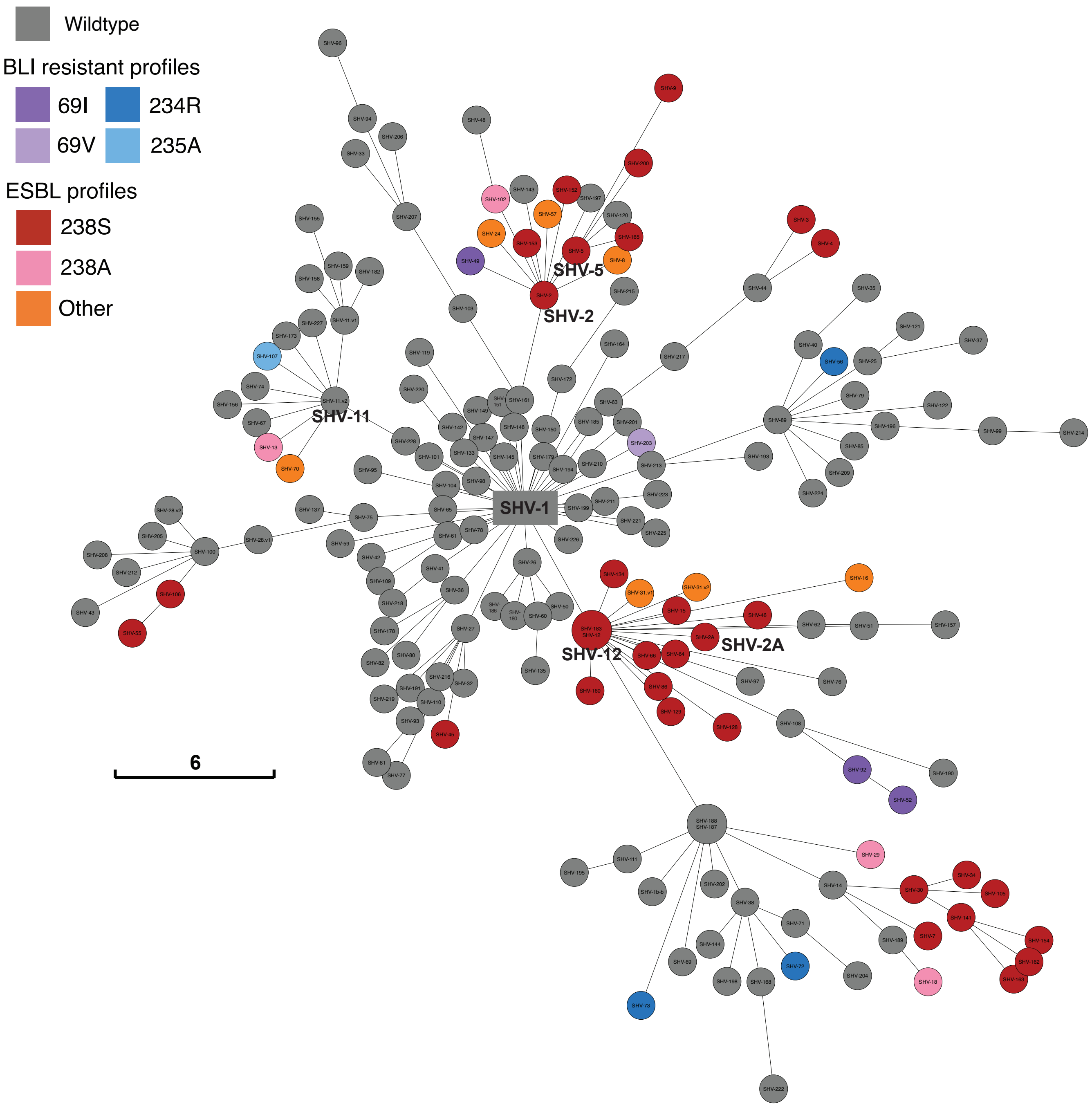

### FigureS2

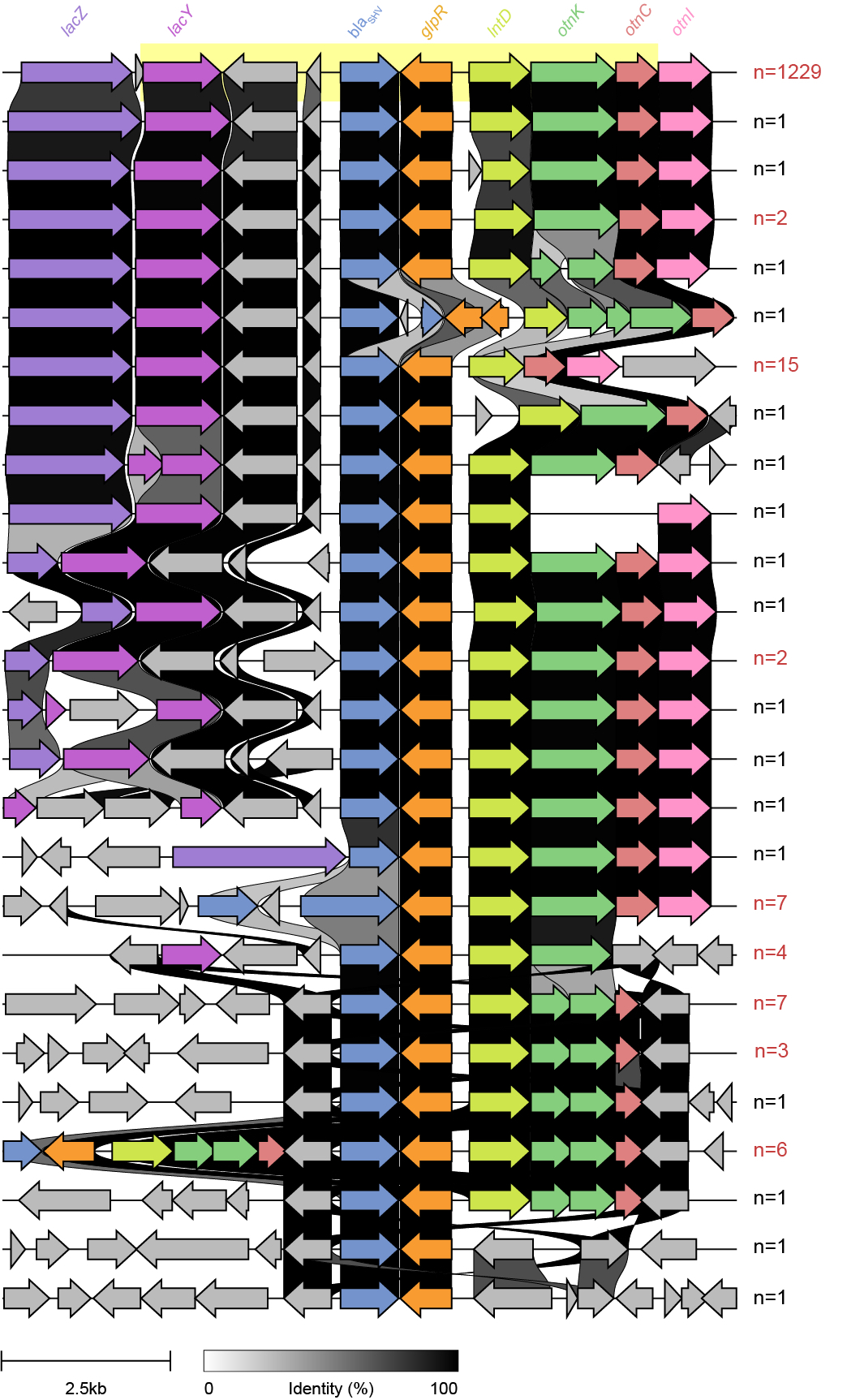
